# An empirical test of the role of small-scale transmission in large-scale disease dynamics

**DOI:** 10.1101/285080

**Authors:** Joseph R. Mihaljevic, Carlos M. Polivka, Constance J. Mehmel, Chentong Li, Vanja Dukic, Greg Dwyer

## Abstract

A key assumption of epidemiological models is that population-scale disease spread is driven by close contact between hosts and pathogens. At larger scales, however, mechanisms such as spatial structure in host and pathogen populations and environmental heterogeneity could alter disease spread. The assumption that small-scale transmission mechanisms are sufficient to explain large-scale infection rates, however, is rarely tested. Here we provide a rigorous test using an insect-baculovirus system. We fit a mathematical model to data from forest-wide epizootics, while constraining the model parameters with data from branch-scale experiments, a difference in spatial scale of four orders of magnitude. This experimentally-constrained model fits the epizootic data well, supporting the role of small-scale transmission, but variability is high. We then compare this model’s performance to an unconstrained model that ignores the experimental data, which serves as a proxy for models with additional mechanisms. The unconstrained model has a superior fit, revealing a higher transmission rate across forests compared to branch-scale estimates. Our study suggests that small-scale transmission is insufficient to explain baculovirus epizootics. Further research is needed to identify the mechanisms that contribute to disease spread across large spatial scales, and synthesizing models and multi-scale data is key to understanding these dynamics.

## Introduction

Ordinary differential equation (ODE) models of host-pathogen interactions rely on the assumption that the host population is well-mixed (Murray, 1989), so that transmission can result from random contact between any given infected/susceptible host pair, with no effects of spatial variation in host density or the environment (Keeling and Rohani, 2008). Such ODE models have led to important conceptual advances, such as the threshold theorem of epidemiology (Kermack and McKendrick, 1927), and the hypothesis that pathogens can control populations of their hosts (Anderson and May, 1979). More recently, the availability of high-performance computing, and the development of sophisticated fitting algorithms, have made it possible to use stochastic versions of ordinary differential equation models, further enhancing their ability to serve as statistical tools for carrying out robust tests of theory (King et al., 2008).

The assumption that pathogen dynamics are driven only by small-scale, spatially-homogenous interactions between individual hosts is perhaps most appropriate for directly transmitted human diseases such as measles and flu (Keeling and Rohani, 2008), and for bite-transmitted animal diseases such as rabies (Blackwood et al., 2013) and facial tumor disease of Tasmanian devils (Hamede et al., 2009). For many other animal diseases, transmission instead occurs when hosts contact infectious pathogen particles in the environment (Rohani et al., 2003), but this complication is often accommodated simply by adding a pathogen-particle equation to otherwise standard models (Anderson and May, 1980). Theory of environmentally transmitted pathogens then follows classical theory in assuming that transmission results from small-scale interactions between hosts and infectious particles, and in assuming that spatial structure and spatial heterogeneity have negligible effects.

For many environmentally transmitted pathogens, however, these assumptions are likely to be incorrect. In bovine spongiform encephalopathy, for example, particle densities in the soil vary spatially (Somerville et al., 2019), while in chronic wasting disease of deer, particle survival in the soil can be altered by spatial variation in soil properties (Kuznetsova et al., 2014). In some *Daphnia* pathogens, infectious particles are ingested during feeding (Shocket et al., 2018), and pathogen dynamics may therefore be modulated by resource quality (Hall et al., 2009), which may in turn vary spatially. In ranaviruses of frogs, transmission rates are partly determined by short-range dispersal of infectious particles (Mihaljevic et al., 2018), which may lead to spatial variation in particle density. In these cases, it seems likely that neglecting spatial structure could lead to deeply flawed model predictions. The reliability of models of environmentally transmitted pathogens is therefore in doubt.

Whether the models are indeed unreliable, however, is unknown, because there are very few tests of the assumption that disease dynamics are driven by contacts between hosts and pathogens at small scales. Part of the problem is that such tests face significant obstacles. Arguably the simplest test would be to compare infection rates at different scales, but data on small scale transmission are often lacking, because epidemiological studies understandably focus on data collected at the scale of the entire host population. Collecting data at both small and large scales could nevertheless provide a robust test of a fundamental model assumption.

A straightforward way to collect infection data at small scales is to carry out transmission experiments. For the vertebrate pathogens that are often the focus of disease ecology, experiments are often impossible (McCallum, 2016), but for some invertebrate pathogens, experiments are possible. For insect baculoviruses, like the baculovirus of the Douglas-fir tussock moth (*Orgyia pseudotsugata*) that we study here, experiments can even be straightforward (Elderd, 2013). In insect baculoviruses, transmission occurs when uninfected host larvae, while feeding on their host plant, accidentally consume infectious particles known as “occlusion bodies”, which are released from the cadavers of dead infected larvae (Cory and Hoover, 2006). For insect baculoviruses, it is therefore possible to carry out experiments on single branches, in which the only process operating is transmission that results from uninfected hosts consuming occlusion bodies released from dead infectious hosts on the same branch.

Because baculoviruses play an important role in controlling pest insects, baculovirus data are also often available at the scale of entire forests (Moreau and Lucarotti, 2007). Forest-scale data are collected to understand the conditions under which natural baculovirus epizootics (epizootics = epidemics in animals) cause the collapse of pest insect populations (Moreau and Lucarotti, 2007), and to document epizootics that result from using baculoviruses as insecticides (Hunter-Fujita et al., 1998). To understand the role of small-scale transmission in baculovirus epizootics, we therefore carried out a small-scale transmission experiment, and we collected large-scale epizootic data from pest control programs in which we participated, and from pest-control programs documented in the literature (Otvos et al., 1987). Our experiment was carried out on single Douglas-fir branches that encompassed an average of 0.15 m^2^ of foliage, while the epizootic data were collected in plots that encompassed 1-10 hectares of forest. The difference in spatial scale over which the two data sets were collected was thus about four orders of magnitude.

To compare pathogen dynamics across spatial scales, we used our short-term, small-scale experimental data to estimate the parameters of a model of baculovirus dynamics, and we inserted the parameters into the model to predict infection rates in epizootics. As we will show, the model is able to explain a substantial fraction of the variation in the epizootic data, but considering only a single model begs the question of whether a model that includes larger-scale processes could better explain the data. Indeed, spatial variation in pathogen densities (Dwyer and Elkinton, 1993) and in forest tree-species composition (Elderd et al., 2013) have been suggested to be key determinant of the dynamics of baculoviruses. A seemingly obvious additional step would therefore be to test whether a model that allows not just for small-scale transmission, but also for large-scale spatial structure or environmental heterogeneity can explain the epizootic data better than a model that allows only for small-scale transmission.

A problem with such an approach is that the epizootic data do not include information about changes in infection rates over space, so it would likely be impossible for us to use the data to make inferences about spatial models. The underlying problem is that the collection of the large-scale data was not guided by an appropriate spatial model. This is a common problem in ecology and epidemiology, especially when theory is confronted by data collected during management programs (Restif et al., 2012).

To avoid this problem, we devised a model-based statistical strategy to infer whether large-scale processes were reflected in the large-scale data, in which we used a proxy model that did not include spatial structure, but that was unconstrained by the small-scale data. This unconstrained model uses the same equations as the model for which we estimated parameters from small-scale data, but its parameters were estimated only from the epizootic data, allowing for the possibility that its parameter values would reflect large-scale spatial variation in host and pathogen densities. To fit both the model with experiment-based parameters and the unconstrained proxy model to the epizootic data, we used Bayesian statistical techniques, which provided a consistent framework with which to compare the two models (Gelman et al., 2014). We thus constructed informative priors for the experiment-based model using the experimental data, and we constructed uninformative or “vague” priors for the proxy model by assuming that all parameters were equally likely, within some large range of possible values.

This approach allows for uncertainty in the experimental parameters, while also allowing for the possibility that spatial structure would lead to differences in the parameter estimates for the two models. As we will show, the parameter estimates are indeed quite different for the two models, and model selection using the Watanabe-Akaike Information Criterion (WAIC, Gelman et al. (2014)) showed that the unconstrained model explains the data far better than the model with experiment-based priors. Interactions between individual hosts and infectious particles at small scales are therefore not sufficient to explain the population-level spread of the tussock moth baculovirus. As we discuss, important missing mechanisms in our models involve large-scale spatial variation, specifically in the frequency of different strains of the baculovirus (Williams et al., 2011), and in the composition of the forests in which tussock moth outbreaks occur (Shepherd et al., 1988). Our results therefore suggest that a better understanding of spatial structure and environmental heterogeneity could significantly improve our understanding of the dynamics of animal diseases, emphasizing the importance of statistically robust empirical tests in the development of ecological theory.

## Methods

### Baculovirus Natural History

The models that we use were first developed for human pathogens (Anderson and May, 1992), but are general enough that they can be used to describe baculovirus epizootics. To explain why, we first describe baculovirus natural history, and then we show how simple SEIR models can encompass this natural history.

Baculovirus epizootics play a key role in terminating tussock moth outbreaks, which occur at roughly 10-year intervals (Mason, 1996). During outbreaks, tussock moth densities increase from levels at which larvae are undetectable, to levels at which defoliation may be widespread and severe (Shepherd et al., 1988). Outbreaking populations are usually terminated by baculovirus epizootics, in which cumulative mortality can exceed 90% (Mason, 1996). So far as can be known, the Douglas-fir tussock moth is the only organism in its range that is susceptible to the baculovirus, although *Orgyia* species from other parts of North America have been successfully infected with the Douglas-fir tussock moth baculovirus in the lab (Rohrmann, 2014).

As is often the case in insect baculoviruses, transmission of the tussock moth baculovirus occurs when larvae accidentally consume infectious virus particles known as “occlusion bodies” while feeding on foliage (Cory and Myers, 2003). Larvae that consume a large enough dose die in about two weeks. Shortly after death, viral enzymes dissolve the insect’s integument, releasing occlusion bodies onto the foliage, where they are available to be consumed by uninfected larvae (Miller, 1997). Epizootics are terminated when larvae pupate, or when epizootics are so severe that most hosts die before pupating (Fuller et al., 2012).

The virus overwinters largely through external contamination of egg masses (Thompson and Scott, 1979). The rate of egg-mass contamination therefore appears to determine infection rates at the beginning of the larval period, which are often low (Otvos et al., 1987). High infection rates then apparently result from multiple rounds of transmission during the larval period (Otvos et al., 1987; Shepherd et al., 1984). In our study areas in Washington, Idaho, and Colorado, USA, and British Columbia, Canada, this period is currently early June to mid-August.

### A Random SEIR Model

Because pathogen transmission and host reproduction occur at different times of the year, we described baculovirus epizootics using a model that does not include host reproduction. We began with a standard Susceptible-Exposed-Infectious-Recovered or “SEIR” model from human epidemiology (Keeling and Rohani, 2008), and we modified the model to allow for two sources of heterogeneity in infection risk. The first source of heterogeneity results from variation between individual hosts, which in insect-baculovirus interactions can be due either to variation in infection risk given exposure to the virus (Páez et al., 2015), or to variation in exposure risk itself (Parker et al., 2010), either of which may be heritable. The second source of variation in infection risk in our model arises from stochastic fluctuations in transmission.

Allowing for stochasticity is important because stochasticity may interact in complex ways with disease dynamics, so that the variability in the model predictions may change as the epizootic proceeds, or as initial host and pathogen densities vary between epizootics. By including stochasticity, we allow for this type of variation, ensuring that our estimation procedures are statistically robust.

Perhaps the best known approach to allowing for stochasticity in ordinary differential equations (ODEs) is to assume that stochastic perturbations occur over infinitesimal time scales, leading to “stochastic ODEs” (Øksendal, 2003). In insect-baculovirus interactions, it seems likely that stochasticity is due to stochastic changes in weather conditions, which can affect baculovirus feeding, and thus infection risk (Eakin et al., 2015). It is therefore intuitive to instead assume that stochasticity operates on a daily time scale.

We thus assume that stochastic perturbations occur over a finite time scale, and so our model equations are known as “random ODEs” (Han and Kloeden, 2017). The distinction from stochastic ODEs is important because it allow us to rely on methods from deterministic calculus (Han and Kloeden, 2017), whereas numerical integration of stochastic ODEs in contrast requires more sophisticated methods (Øksendal, 2003).

Our approach is to first construct a model for epizootic dynamics during a single day:

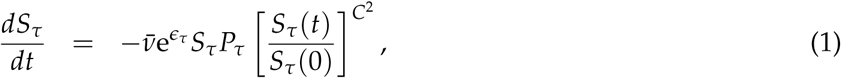

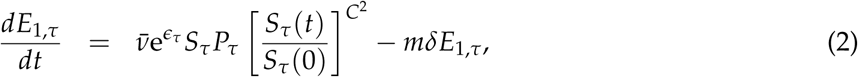

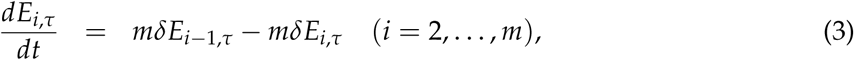

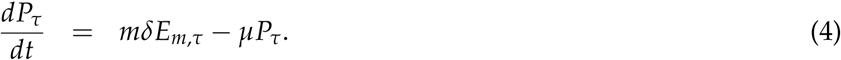

Here the subscript *τ* is an integer denoting the day, so that the stochasticity term *ϵ_τ_* represents the stochastic perturbation on day *τ*. In the interests of simplicity, we assume that *ϵ_τ_* follows a normal distribution with mean 0 and standard deviation *σ*, and we exponentiate *ϵ_τ_* to avoid negative transmission rates, which would be biologically nonsensical. Because weather conditions likely varied between populations, we estimated a value of the standard deviation *σ* for each epizootic.

As in standard SEIR models, transmission in this model occurs through a mass action term. In models of human diseases, this term is often written as *βSI*, where *S* is the density of uninfected or “susceptible” hosts, *I* is the density of infected hosts, and *β* is the transmission parameter. Here transmission instead occurs through contact between susceptible hosts *S* and infectious cadavers *P*, with transmission parameter 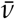.

We further modify the transmission term to allow for inherent variation among hosts, according to 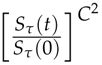. This term comes from models originally developed for the gypsy moth baculovirus (Dwyer et al., 1997), which were in turn derived from models of sexually transmitted infections in humans (Anderson and May, 1992). The approach is to assume that there is a distribution of infection risk in the host, with mean 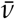 and coefficient of variation *C* (Dwyer et al., 2000), as an approximation to a more computationally intensive partial-differential equation model. The approximation is highly accurate if transmission rates follow a gamma distribution, but it is also reasonably accurate if transmission rates instead follow a log-normal distribution, which has a fatter tail than a gamma distribution (G. Dwyer, unpublished). In general, we assume that transmission occurs among larvae in the fourth instar (= larval stage), the instar that has the biggest impact on cumulative infection rates (Dwyer, 1991; Otvos et al., 1987). To allow for the smaller size of hatchlings, we multiply the initial virus density by the parameter, *ρ*, which is the ratio of the number of virus particles produced by a first-instar cadaver to the number produced by a fourth-instar cadaver.

As in standard SEIR models, susceptible hosts *S_τ_* that become infected proceed through *m* exposed classes, *E_i_*_,*τ*_. Exposed hosts eventually die of the infection, joining the infectious-cadaver class *P_τ_*, which represents the dynamics of the pathogen in the environment (the *R* class of SEIR models corresponds to cadavers that are no longer infectious, and so we do not include it here). Pathogen infectiousness decays at rate *µ*, due mostly to UV radiation (Thompson and Scott, 1979). Hosts move between exposed classes at rate *mδ*, so that the time spent in a single exposed class follows an exponential distribution with mean time 1/(*mδ*). The total time in the *m* exposed classes is the sum of *m* such distributions, and a well-known theorem has shown that this sum follows a gamma distribution with mean, 1/*δ*, and variance, 1/(*mδ*^2^) (Keeling and Rohani, 2008).

Once equations (1)-(4) have been numerically integrated for day *τ*, the initial conditions for day *τ* + 1 are updated. To carry out this updating, the model sets the initial conditions for day *τ* + 1 equal to the final values on day *τ*:

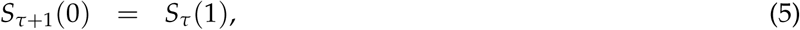

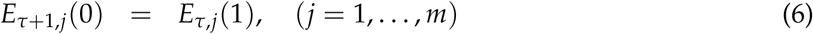

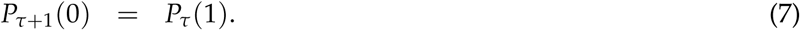

Here *S_τ_* (1) is the density of uninfected larvae at the end of day *τ*, while *S_τ_*_+1_(0) is the density of uninfected larvae at the beginning of the following day, and so on for the other state variables.

### Field transmission experiments

In equations (1)-(4), the average transmission rate, 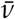, represents the overall infection risk per unit time. It therefore encompasses both the probability of exposure and the probability of infection given exposure, and so it allows for the effects of both host behavior and host-tree foliage quality. Transmission 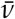 is therefore best measured in the field, so that larvae can feed freely on virus-contaminated foliage (Elderd, 2013).

Previous work with the gypsy moth, *Lymantria dispar*, produced a protocol for baculovirus field-transmission experiments that gave repeatable results (Dwyer, 1991; Dwyer et al., 1997; Fleming-Davies et al., 2015). Following this protocol, we first reared uninfected tussock moth larvae in the laboratory from field-collected egg masses. The egg masses had been collected from an early-stage outbreak in Cheyenne Mountain State Park in Colorado, USA, in 2014. To deactivate any virus particles on the surface of the eggs, we submerged the egg masses in 5% formalin for 90 minutes prior to incubation (Dwyer and Elkinton, 1995). We used hatchling larvae as infected hosts in our experiments because the most important round of transmission occurs when third and fourth instar larvae are infected by first instar cadavers (Otvos et al., 1987; Shepherd et al., 1984).

To infect the hatchlings, we placed them on artificial diet contaminated with the virus. A pilot study allowed us to determine the viral dose that results in roughly 95% of larvae becoming infected. We therefore used a solution of 10^4^ occlusion bodies/*µ*l, and we used a Pasteur pipette to place 5 drops of this solution onto artificial diet in a 6 oz (177 ml) plastic rearing cup. After larvae fed for 24h, they were moved to additional rearing cups.

To ensure that the infected larvae were indeed infected, we reared them at 26 °C in the laboratory for five days. Five days was long enough to ensure that any uninfected larvae would molt to the second instar, whereas the virus prevented infected larvae from molting (Burand and Park, 1992). Second instars have different coloring from first instars, and so it was straightforward to identify and remove uninfected larvae, which were all in the second instar (Fuller et al., 2012). The infected larvae were then placed on Douglas-fir branches in the field at two densities, 10 and 40 larvae per branch. The trees that we used were encompassed within an area of roughly 2 hectares in the Okanogan-Wenatchee National Forest, near Entiat, Washington, USA.

The branches were enclosed in mesh bags, which prevent the emigration of larvae and the breakdown of the virus (Fuller et al., 2012). We then allowed 5 days for the infected larvae to disperse on the foliage and die, a time sufficient to ensure that they all died. On the fifth day, we added 20 uninfected larvae, which we had reared in the lab to the fourth instar (Dwyer et al., 1997). Controls consisted of branches containing only uninfected larvae.

An insect larva’s susceptibility to baculovirus infection can vary in a complex way within an instar (Grove and Hoover, 2007). We therefore developmentally synchronized the uninfected fourth instars, as follows. Shortly before molting, a larva’s head capsule slips forward, making it possible to see that the larva is close to the end of its instar. To synchronize fourth instars, we collected third instars with slipped head capsules, and we held them at 4°C, halting development, until we had enough larvae to begin the experiment. We then reared the larvae at 26°C until they had molted to the fourth instar, which occurred within 48 hours. The effect was that the uninfected larvae all reached the fourth instar within a relatively short period. In previous experiments that used this protocol, the variance in the infection rate was indistinguishable from the variance of the corresponding binomial distribution (Elderd et al., 2008). Synchronization therefore appears to eliminate most sources of extraneous variability, leaving only the binomial variation that is expected in an infection experiment.

Our experimental treatments consisted of the 2 viral densities (10, 40), crossed with 3 viral isolates, for a total of 6 treatments, each replicated 14 times. We also had 7 control bags, in which there were no infectious cadavers. The WA isolate was collected from Washington State in 2010, while the NM isolate was collected in New Mexico in 2014. The final isolate was TMB-1 (“Tussock Moth Bio-control 1”), which makes up the insecticidal spray formulations produced by the USDA Forest Service (Martignoni, 1999). Transmission electron microscopy showed that all three isolates were the multicapsid form of the virus, known as “*Op*MNPV”. In multi-capsid strains, the virions are found in clumps within the occlusion bodies, as opposed to single-capsid (*Op*SNPV) strains, in which the virions occur singly within the occlusion bodies (Hughes and Addison, 1970).

The experiment included 91 branches with 20 uninfected larvae each, for a total of 1,820 uninfected larvae. We allowed the initially uninfected larvae to feed on foliage for 7 days, and then we removed the branches from the trees and brought them into the laboratory. Next, larvae from the branches were reared individually for three weeks in 2 ounce (59 ml) cups partially filled with artificial tussock moth diet, at 26 °C in the laboratory. To determine if larvae had died of the virus, we examined smears from dead larvae under a light microscope at 400*×* for the presence of occlusion bodies, which are easily visible at that magnification (Fleming-Davies et al., 2015).

After the 7 day experimental period, we photographed each branch, and we used ImageJ (Schindelin et al., 2015) to estimate the area of foliage on each branch. We then calculated the density of infectious cadavers and the density of uninfected hosts by dividing the number of cadavers or hosts by the foliage area. This allowed us to measure densities on single branches using the same units as in the epizootic data, with the proviso that the epizootic data were collected at a much larger scale.

As part of this experiment, we attempted to measure the decay rate *µ* of infectious cadavers on foliage, by allowing some cadavers to be exposed to sunlight outside of the mesh bags for 3 days. 3 days, however, proved to be too short a period for us to detect meaningful decay of the virus, and we therefore do not report those results. Previous work was similarly unsuccessful at estimating the decay rate of the tussock moth baculovirus using exposure periods of 1, 4, 13, and 32 days (Dwyer, 1992). It may therefore be that the decay rate of the virus is very low. As we will show, this result is consistent with some of the estimates of the decay rate *µ* that result from fitting our models to the epizootic data.

#### Estimating Transmission Parameters 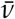 & C From the Experimental Data

Direct comparison of the infection rate in our experiment to the infection rate in the epizootics is not meaningful, because our experiments were designed to only allow for a single round of transmission, whereas there were undoubtedly multiple rounds of transmission in the epizootics. The infection rate in the experiment was therefore likely to differ from the infection rate in the epizootics simply because of differences in time scales, rather than because of differences in spatial scales.

To correct for the difference in time scales, we fit a simplified version of our SEIR model to the experimental data, to estimate the transmission rate in the experiment. We then used the resulting estimates of the average transmission rate 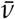 and the heterogeneity in transmission parameter *C* in the full model, and we compared the model predictions to the epizootic data, as we described. This approach allowed us to compare transmission rates at the two spatial scales in a way that corrected for the difference in temporal scales.

To simplify the full model, we first assumed that transmission stochasticity was negligible during the short time scale of the experiment. This allowed us to eliminate the dependence on *τ* in equations (1)-(4). Also, the mesh bags that enclosed the experimental branches prevented the emigration of larvae and the breakdown of the virus, and the experiment was short enough that no larvae became infected and died during the experiment. The density of infectious cadavers on the experimental branches was therefore constant after the experiment began, so the density of particles *P* was constant during the experiment. This latter simplification allowed us to solve equation (1) for the fraction of hosts *i* that have become infected by the end of the experiment, as a function of the initial (and constant) virus density *P*_0_ (Dwyer et al., 1997).

When we compare the model to the experimental data, it is useful to write the expression for the fraction infected in terms of the log of the fraction uninfected (1 *− i*):

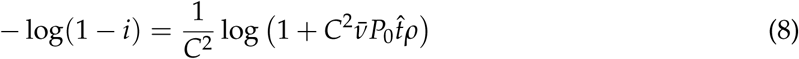

Here, 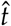 is the time larvae were exposed to virus on foliage, which was 7 days. The ratio parameter, *ρ*, is included because an implicit assumption of the SEIR model is that all larvae are in the fourth instar, but the infectious cadavers in our experiment were in the first i nstar. T he p arameter *ρ* therefore scales the transmission rate so that it is expressed in terms of fourth-instar cadavers. We also considered a model in which host heterogeneity *C* is negligible:

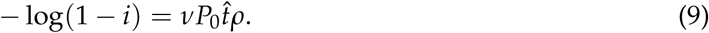

To estimate average transmission 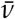 and heterogeneity *C* from the experimental data, we used the Bayesian inference software, *JAGS* (http://mcmc-jags.sourceforge.net/) via the *rjags* package in R. We assumed vague priors for 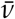 and *C*, and an experimentally-derived prior for the ratio parameter *ρ*, as we will describe. To avoid biases in the parameter estimates, we explicitly allowed for error in the cadaver densities *P*_0_ in the statistical model. We then used WAIC to compare the ability of different models to explain the data (see Online Appendix for the definition of WAIC). Example code for this fitting routine is in the Online Appendices.

On seven of our 91 experimental branches, all the initially uninfected larvae died due to desiccation. Desiccation is a common source of natural mortality in Douglas-fir tussock moth populations (Mason and Torgersen, 1983), but it may have been slightly worse in our case because the mesh bags can elevate temperatures (Páez et al., 2017). We therefore excluded these seven branches from our analyses. We also excluded desiccated larvae from the data from the other branches, partly because desiccated larvae were too dry to be autopsied, but more importantly because desiccated larvae were unlikely to have been infected.

Because the transmission rate 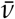 is scale dependent, we measured the foliage area of the branches in our experiment. This area ranged from 0.09 to 0.28 *m*^2^ (mean = 0.15). After we corrected for branch area, the cadaver densities for the low density treatment (10 cadavers per branch) ranged from 38.49 to 112.58 cadavers per m^2^ of foliage (mean = 70.58), while the densities for the high density treatment (40 cadavers per branch) ranged from 142.38 to 426.98 cadavers per *m*^2^ of foliage (mean = 254.41).

#### Estimating Speed of Kill 1/δ and Ratio ρ From Experimental Data

We also used an experiment to estimate the average speed of kill, which in the model is equivalent to 1/*δ*, the inverse of the death rate. In this experiment, we infected larvae by allowing them to feed on Douglas-fir foliage that was contaminated with a sprayed virus solution (Online Appendices). For logistic convenience, this experiment was carried out in the laboratory. Because speed of kill is affected by temperature, we held the larvae at temperatures typical of field conditions (Polivka et al., 2017).

To estimate the ratio parameter, *ρ*, we again infected larvae in the laboratory, but in this case we infected both hatchlings and fourth instars (Online Appendices). We held these larvae in the laboratory until death or pupation, and we counted the number of occlusion bodies per dead larva for each instar, using a hemocytometer under a light microscope.

### The Epizootic Data

Our epizootic data came both from naturally occurring epizootics, and from pest management programs that used the virus as an environmentally benign insecticide to reduce tussock moth defoliation. Our data set included 7 unsprayed control plots and 5 sprayed treatment plots, with data coming both from the literature (Otvos et al., 1987), and from data that we present here for the first time. Although it is at least possible that the TMB-1 isolate used in the spray formulation is phenotypically different than wild-type virus, fitting models with different transmission rates for spray and control plots showed that the transmission rates of sprayed and wild-type virus are effectively indistinguishable.

In spray programs, managers typically establish control plots in the same general area as spray plots, but far enough away to prevent sprayed virus from drifting into the controls. Data from Washington State, for example, were from a spray program in 2010, in which all plots were at least 10 kilometers apart. This distance is far enough that drift of the virus spray was highly unlikely. The data from British Columbia were similarly from a spray program in 1982 (Otvos et al., 1987), but some plots were only a few hundred meters apart. In the control plots in British Columbia, however, the epizootics started 3-4 weeks later than in the treatment plots. Given that sprayed virus typically decays within a few days (Polivka et al., 2017), this time lag suggests that drift of the spray was again minimal.

At the beginning of the larval period at each site, initial host and pathogen population densities were estimated using standard methods. These initial densities provided initial conditions for the model (Online Appendices). Because initial virus densities in sprayed treatment plots were much higher than in unsprayed control plots, the two types of epizootic data together encompass a broader range of initial pathogen densities than either type of epizootic data alone. This is important because a broad range of densities often increases statistical power when ecological models are fit to data (Pascual and Kareiva, 1996).

The data then consist of the fraction of larvae infected, estimated at intervals of roughly a week, for up to 50 days, typically from mid-June to mid-August. In the sprayed plots, insects were collected within 7 days of the application of the virus, but in the control plots the start of collections was more variable, particularly at sites where there was no concurrent spray project. Insects were reared and diagnosed as in the field transmission experiment.

#### Model fitting and model selection

To fit models to the epizootic data, we compared the fraction infected in the data to the fraction infected in the model. In the model, infected (but not yet dead) larvae are represented by the exposed classes *E_m_*. The fraction of larvae infected is then 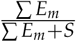.

For the transmission data, we used a binomial likelihood function, but for the epizootic data, a binomial likelihood function was unlikely to be sufficient. Use of the binomial distribution rests on the assumption that individual hosts are independent (McCullagh and Nelder, 1989), which likely held in our field experiment, as in similar field experiments (Elderd et al., 2008), but the environment in which epizootics occur is much more complicated. For example, the density of hosts may have been clumped within the forest, and this clumping could cause the variance in the infection risk to be substantially higher than the variance of the corresponding binomial, a phenomenon known as over-dispersion. It was therefore important to allow for the possibility of over-dispersion.

In the absence of direct information on the level of over-dispersion, a useful approach is to use a beta-binomial distribution (Cox and Snell, 1989). In a beta-binomial, the binomial probability of an infection *p* follows a beta distribution, which describes quantities like *p* that vary between 0 and 1. The beta-binomial then has two parameters, as opposed to the single parameter of the binomial, making it possible to increase the variance of the likelihood, as needed, to explain the lack of fit of the model to the data (by including stochasticity in transmission, we also allowed for the possibility that the lack of fit was due to stochasticity). As parameters of the beta-binomial, we used *a* = *pe^γ^* and *b* = (1 *− p*)*e^γ^*, where *p* is the model prediction of the fraction infected and *γ* is an inverse measure of the over-dispersion. As we will show, over-dispersion levels were moderate but not excessive.

Because our epizootic models allow for stochastic fluctuations in transmission, we integrated values of the likelihood over many realizations of the models. This approach is equivalent to integrating out the values of the stochasticity *ϵ_τ_* to produce an average likelihood:

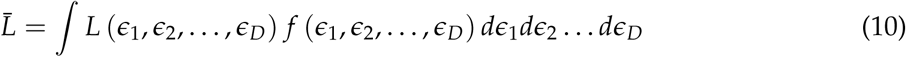

Here 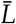 is the average likelihood, and *D* is the number of days in an epizootic. *D* is thus the number of days for which we drew values of *ϵ_τ_*. The function *f* (*ϵ*_1_, *ϵ*_2_, …, *ϵ_D_*) is the probability density of the *ϵ_τ_*’s, where each integer, 1, 2, …, *D*, indicates a different day.

Numerical integration of the model is computationally expensive, and using numerical quadrature to calculate the integral in equation (10) is therefore impractical. Accordingly, we instead used Monte Carlo integration (Ross, 2002). This meant that we drew values of the *ϵ_τ_*’s, and then we estimated the average likelihood, according to;

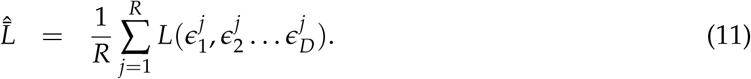

Here *R* is the number of realizations, and 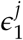 is the value of *ϵ*_1_, meaning the stochastic term on day 1, in the *j*th realization, and so on for 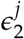 … 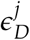. According to the weak law of large numbers, as 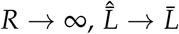 (Ross, 2002).

To reduce computing time, it was important to ensure that 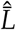 approached 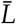 for a reasonably small number of realizations. For this purpose we used the MISER Monte-Carlo integration algorithm. This algorithm uses recursive, stratified sampling to estimate the average likelihood, while minimizing the number of realizations. Briefly, the algorithm works as follows (the code that we use is from the Gnu Scientific Library, but for a clear explanation of how the algorithm works, see Press et al. (1992)). As equation (11) shows, in calculating an estimate of the average likelihood 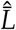, we are sampling over a *D*-dimensional space of stochasticity parameters *ϵ_τ_*. Within this parameter space, it is likely that there are sub-spaces within which the variance in 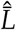 is higher than in other sub-spaces. In estimating 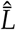 across the entire space, it turns out to be more efficient to sample more frequently in sub-spaces within which the variance is higher. There is a formal proof of this proposition, and so the process of sub-sampling forms the basis of the MISER algorithm, as follows.

The algorithm is given a quota of *R* realizations. Some fraction of these realizations, in our case 0.1, is devoted to sampling uniformly across the entire space. Based on this initial sample, the algorithm recursively divides the overall sample space into sub-spaces of high and low variance. In using the remaining realizations, the algorithm samples more intensively in sub-spaces of high variance. The end result is an estimate of 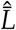 that minimizes the variance. Initial trials with this algorithm showed that 150 realizations was usually sufficient to produce reliable estimates of 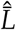. Using a larger number of realizations only reduced the variance in 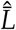 by a small amount.

We then used our likelihood in Bayes’ theorem:

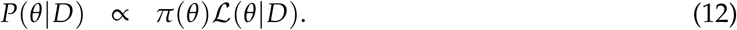

Here *P*(*θ|D*) is the posterior probability distribution of the parameters *θ* of our model, which we fit to the data *D*. The symbol *π*(*θ*) is the prior probability of the parameters, and *L*(*θ|D*) is the likelihood of the parameters. Because posterior probabilities are generally only used for comparison purposes, we only need to calculate the posterior probability up to a constant of proportionality, and so we use the proportion symbol ∝.

To create priors from our experimental data, we used the data to construct log-normal prior distributions for transmission 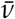, heterogeneity *C*, and ratio *ρ*. To do this, we used the marginal posterior samples generated from fitting the model to our transmission data. On a log scale, the marginal posterior samples were well described by normal distributions, so we used log-normal priors with means and standard deviations calculated from the posterior samples. In the case of the heterogeneity parameter *C*, we instead estimated *k* = 1/*C*^2^, because *k* has a distribution that is closer to normal than *C*, after being log-transformed.

We similarly used our speed of kill data to construct a log-normal prior on the death-rate parameter *δ*. Because, in the model, the variance in the speed of kill is determined by the number of exposed classes *m*, in principle it should be possible to estimate *m* from the observed variance in the speed of kill in experimental data. In practice, however, the variance in the speed of kill in experimental data is usually very low (Dwyer, 1991). Meanwhile, our preliminary efforts to estimate *m* from the epizootic data were unsuccessful. Accordingly, instead of estimating *m*, we set *m* = 200. Fixing *m* at this value ensured that the variance in the speed of kill was realistically low, without the necessity of estimating the uncertainty in *m*.

The model parameters that we fit to the epizootic data were the decay rate *µ*, the stochasticity parameter *σ*, and the over-dispersion parameter *γ*. For these latter parameters, we used uniform probability distributions as “vague” priors, so that effectively all possible parameter values were equally likely, up to some high upper limit. The parameters *σ* and *γ* in particular determine the process error and the observation error respectively (Bolker, 2008), and are thus effectively “nuisance” parameters. The only biologically interesting parameter that was unconstrained by the epizootic data was therefore the cadaver decay rate *µ*.

By using Bayes’s theorem, we allowed for the possibility that the likelihood would dominate the experiment-based priors, and it was therefore possible that our fitting routine would produce posterior parameter estimates that were far from the values calculated from our experiments. For the model with experiment-based priors, however, the posterior median values of the parameters were not that far from the medians calculated from our experiments. This could have happened because the experiment-based priors provide an excellent fit to the epizootic data, but it could also have happened because the epizootic data did not provide much information about the model and its parameters. In practice, both phenomena were operating, in the sense that the experiment-based priors provide a reasonable fit, and that the epizootic data provided only a moderate amount of information about the model parameters.

To show this, we compare the parameter values for the model with experiment-based priors to a model in which the corresponding parameters have vague priors. Differences in the posterior distributions of the parameters for the two models then indicate first that the experimental data did indeed constrain the posterior estimates of the parameters for the model with experiment-based priors. As we will show, for the heterogeneity parameter *C*, the ratio parameter *ρ*, and the death-rate parameter *δ*, the posterior medians for the model with vague priors were close to the posterior medians for the model with experiment-based priors, but the posterior median of the transmission rate 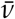 was meaningfully different from the median for the model with experiment-based priors.

It is also important to remember that the model with all-vague priors is the model that we use as a proxy for more complex models that take into account processes above and beyond the processes that take place on a single branch. From this perspective, the difference in posterior median transmission rates between the model with experiment-based priors and the model with all-vague priors is important because the transmission rate 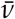 reflects the scale at which interactions occur. Partly for this reason, 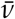 is in units of per infectious cadaver per m^2^, per day.

To compare the fit of the model with all-vague priors to the fit of the model with experiment-based priors, we first calculated the coefficient of determination *r*^2^ for each model. To define *r*^2^, we first define SS_tot_ to be the total sum of squared errors across all observations in our data set:

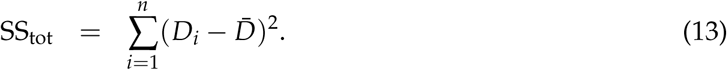

Here *n* is the total number of observations of the fraction infected in the epizootic data, *D_i_* is data point *i*, and 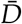 is the average fraction infected across all epizootics. SS_tot_ is thus the total variation in the data set. Also, SS_res_ is the residual sum of squares, defined as:

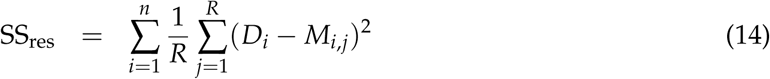

Here we are averaging across *R* = 500 model realizations. SS_res_ thus measures the error between the model and the data, which is the extent to which the model reproduces the data. We then define *r*^2^ according to,

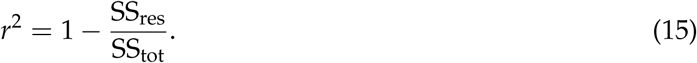

We thus use *r*^2^ to calculate the fraction of the variance in the data that is explained by the model, or alternatively, the extent to which the model produces better predictions of the epizootic data than a simple prediction that the fraction infected at each time point in each population is equal to the average fraction infected across the entire dataset.

Because the model with vague priors was fit only to the epizootic data, its *r*^2^ value was guaranteed to be as good or better than the *r*^2^ value for the model with experiment-based priors. As we mentioned, however, the improvement in the *r*^2^ value turned out to be modest. An additional important question is therefore, how much better is the fit of the model with vague priors than the fit of the model with experiment-based priors? That is, does the model with vague priors provide a meaningfully better explanation for the data than the model with experiment-based priors? To consider which processes in particular are poorly described by our experiments, we also considered models that allowed for experiment-based priors on only some of the parameters for which we had experimental data.

We then used statistical model selection to compare the ability of the different models to explain the epizootic data. Because Bayesian statistical techniques are fundamental to our approach, we chose between models using the Watanabe-Akaike Information Criterion or WAIC, a Bayesian version of the more familiar AIC (Gelman et al., 2014). In most applications of model selection, the models being compared differ in structure, but in our case, the model structure, as defined by the random ODEs in equations (1)-(4), is the same for all models. Because WAIC is a type of Bayesian information criterion, it allowed us to choose between models that differed only in their prior probability distributions. We are thus carrying out model selection in an unconventional way, but to our knowledge there is no established method of choosing between models with different priors. This is true even though estimating model parameters at a smaller scale than the test data is a common procedure in disease ecology. Using small-scale data to construct priors is one way to estimate model parameters at a smaller scale than the test data, in a way that allows for parameter uncertainty (Elderd et al., 2006). We therefore argue that WAIC is useful for testing whether small-scale data can explain large-scale data.

As we will show, the results of our WAIC analysis confirm the results of our comparisons of posterior parameter estimates. That is, models with experiment-based priors on the transmission parameter fit the epizootic data substantially worse than models with vague priors on the transmission parameter. We therefore conclude that the dynamics of the baculovirus are partly affected by processes at larger scales than the scale at which individual hosts interact.

## Results

### Experiments

For two of the three virus strains that we tested, the overall best-fit model is clearly nonlinear (fig. 1). The model with the best (lowest) overall WAIC score therefore includes host heterogeneity in infection risk (Table 2). Fig. 1 also shows that the three isolates differed strongly, such that the NM isolate had much higher heterogeneity in transmission, while the WA isolate had much lower heterogeneity in transmission. These effects are reflected in the median posterior estimates of heterogeneity *C* for the three isolates (Table 1).

**Figure 1:**
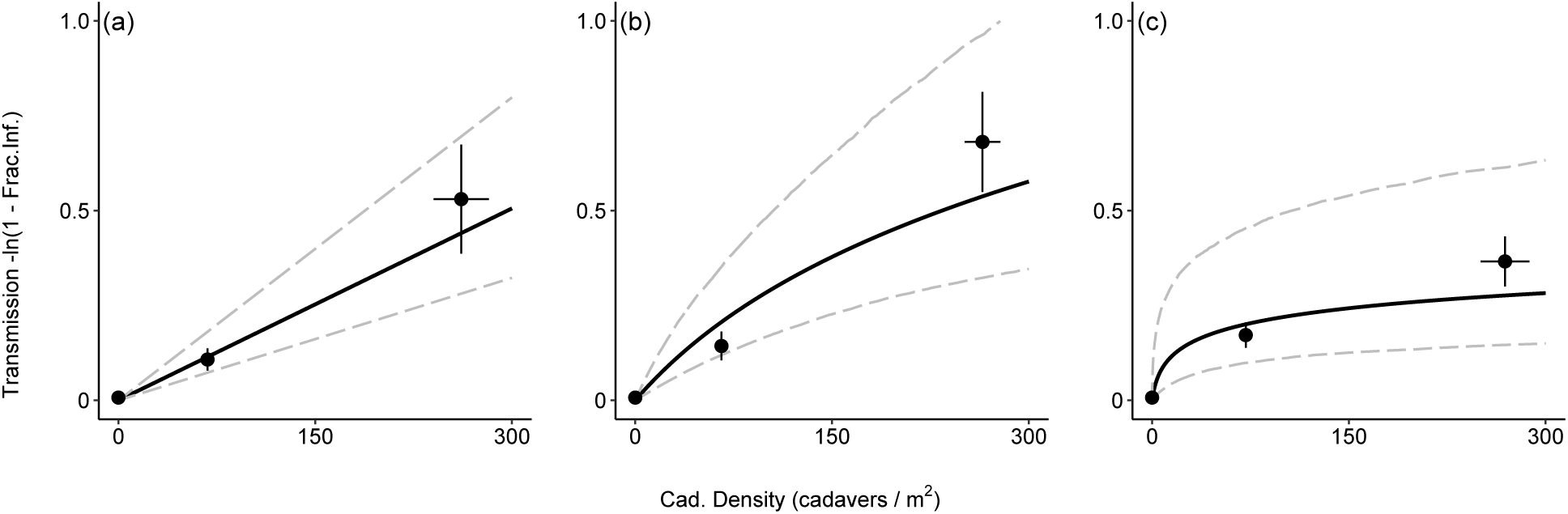
Results of the field transmission experiment, with (a) the WA isolate, (b) TMB-1, and (c) the NM isolate. In the figure, the solid lines represent the median model predictions, while the gray dashed lines represent boot-strapped 95% credible intervals. The large black points with error bars represent the mean and the standard error of the data.

**Table 1:** Best-fit transmission parameters (average transmission, 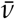, and heterogeneity, *C*) for three viral isolates. Values are posterior medians with 95% credible intervals. Units on 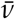 are per infected cadaver, per m^2^, per day. Heterogeneity is the squared C.V. of the distribution of infection risk, and it is therefore scale free.

| Isolate | Average transmission, $\bar{v}$ | Host heterogeneity, $C$ |
| --- | --- | --- |
| WA | 0.006 (0.003, 0.010) | 0.60 (0.05, 1.52) |
| TMB-1 | 0.012 (0.004, 0.030) | 1.52 (0.25, 2.51) |
| NM | 0.076 (0.005, 3.104) | 4.11 (2.15, 6.45) |

**Table 2:** Model selection for the transmission experiment. The bold-faced WAIC scores highlight the best model, based on ΔWAIC > 3.

| Model Type | WA Isolate | NM Isolate | TMB-1 | Overall |
| --- | --- | --- | --- | --- |
| $C > 0$ | 175.53 | **179.48** | **165.06** | **520.07** |
| $C = 0$, | 174.28 | 199.11 | 168.45 | 541.84 |

In the gypsy moth baculovirus, there is a negative correlation between the average and the CV of transmission (Fleming-Davies et al., 2015). Table 1 at least suggests that such a correlation may similarly occur in the tussock moth baculovirus, but with only 3 isolates, we cannot reach general conclusions. More immediately, the variation across isolates is important because, in the epizootic data, we have no information about the isolates that were present. In constructing informative priors from our experimental data, we therefore allowed for variation across isolates by pooling the marginal posterior distributions for the three isolates, and inflating the pooled variance slightly (Online Appendices).

### Comparing Models to the Epizootic Data

In the epizootic data, initial infection rates in control populations were low, but increased slowly over the larval period (fig. 2), taking weeks to reach high levels. Sprayed populations in contrast received an initial inundation of the pathogen, and so their infection rates increased within a week or two after the spray application. Host population collapse was therefore rapid in the sprayed sites, leading to weaker effects of initial host density, as opposed to initial virus density.

**Figure 2:**
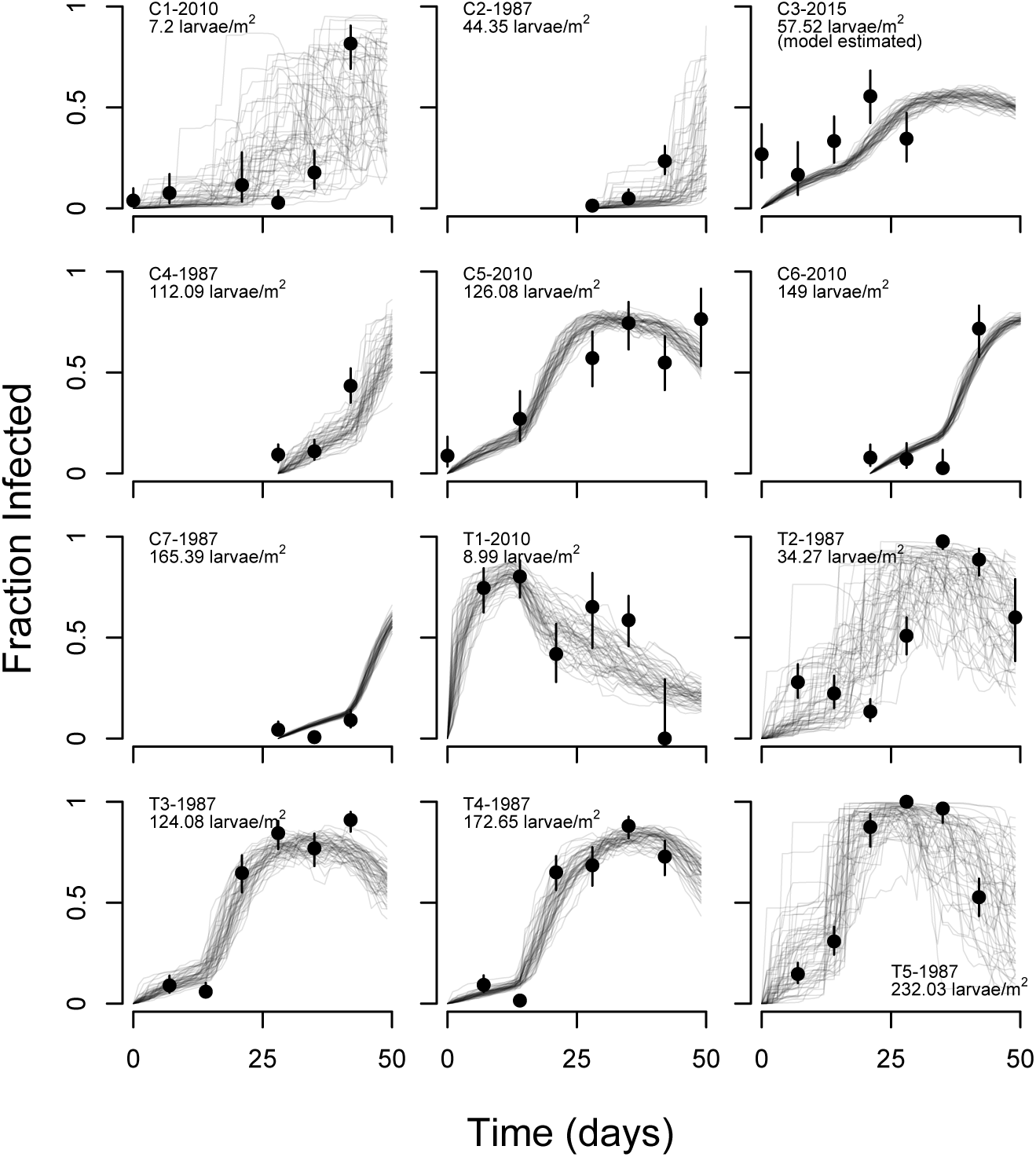
Stochastic realizations of the model with informative priors on 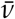, *C*, *ρ*, and *δ*, versus the data (black points with 95% binomial confidence intervals as error bars). The labels C and T stand for Control and Treatment (meaning treated with virus spray), respectively, and are followed by the year of observation. The initial larval host density is also shown. Note that for the Colorado site (C3-2015), the initial larval host density was estimated from the data. Here and in subsequent figures, for two of the populations (C3-2015, C4-1987), we show the model’s predictions after the last data point was collected, to illustrate the overall dynamics of the pathogen.

The model with experiment-based priors generally does a good job of reproducing the epizootic data (fig. 2), with *r*^2^ = 0.68, while the model with vague priors improves on the fit only modestly (fig. 3), with *r*^2^ = 0.75. The posterior estimates of the model parameters, however, show that the models provide quite different explanations for the dynamics of epizootics.

**Figure 3:**
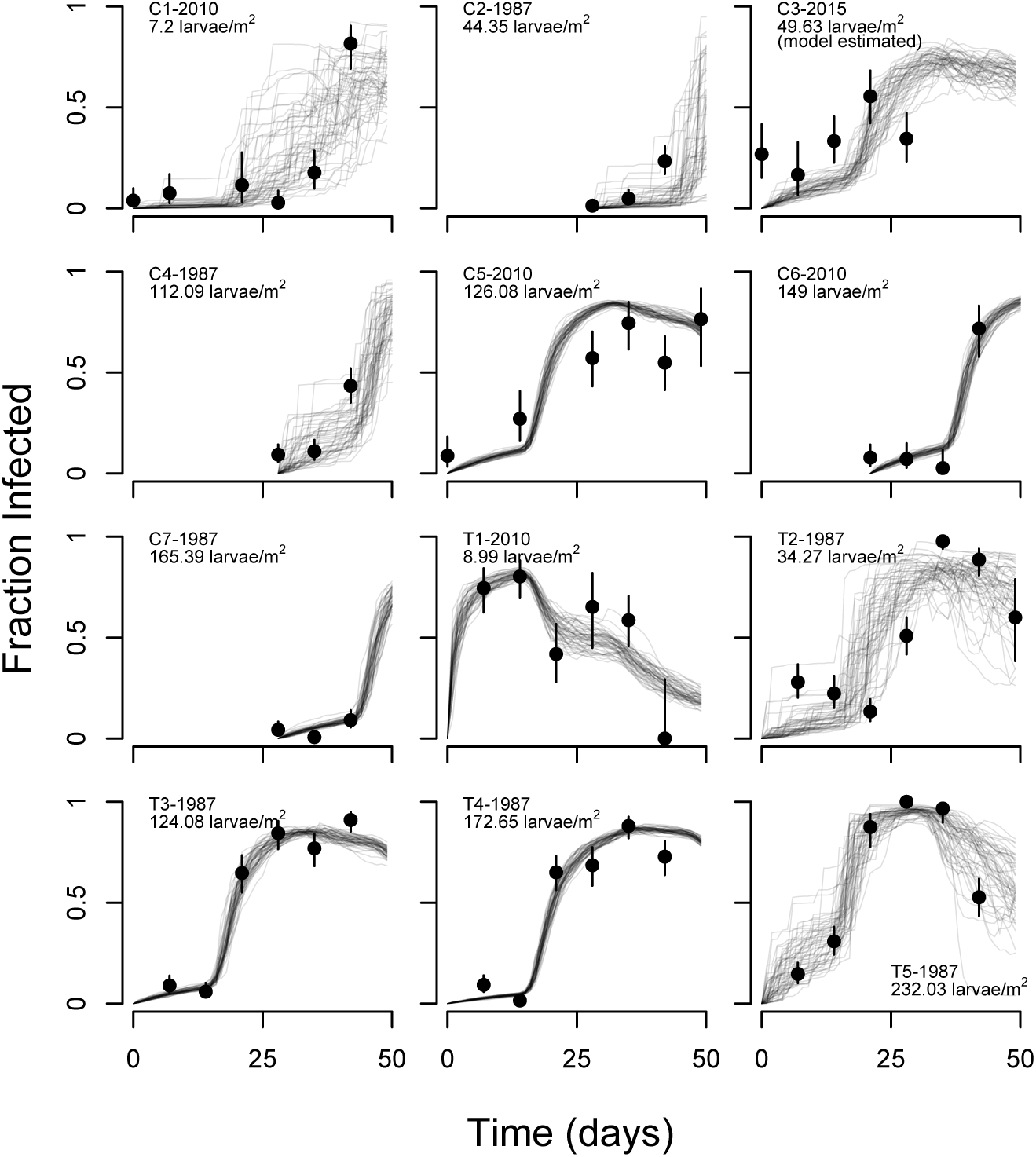
Stochastic realizations of the model that uses vague priors on all parameters, with symbols and labels as in fig. 2.

For both models, the posterior distributions of the heterogeneity parameter *C*, the ratio parameter *ρ*, and the speed of kill parameter *δ* strongly overlap with the posteriors from the experimental data (fig. 4). For models with intermediate numbers of experiment-based priors, the posterior distributions of these three parameters also strongly overlap with the experimental posteriors (fig. 5). These results suggest that our experimental estimates of heterogeneity *C*, the ratio parameter *ρ*, and the death rate parameter *δ* are all reasonably accurate, compared to estimates that take into account epizootic data.

**Figure 4:**
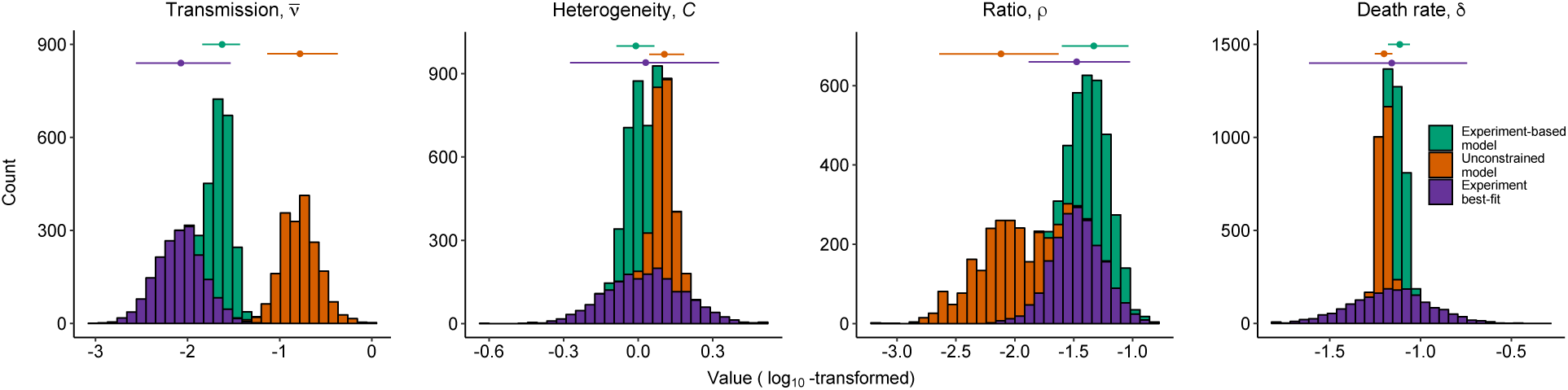
Comparison of posterior distributions for the 4 model parameters for which we have small-scale experimental estimates: the average transmission rate 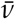, the heterogeneity in transmission *C*, the ratio parameter *ρ*, and the death rate parameter *δ*. For each parameter, we compare the posterior marginal distribution from our experimental data (purple), to the posterior marginals when we fit models to the epizootic data. The latter distributions include the case in which we used experiment-based priors (green), and the case in which we used uninformed priors (orange). Points with error bars near the top of each panel show the median and the 95% credible interval in each case. Note that each panel has a different scale on its horizontal axis.

For both the unconstrained model and the model with experiment-based priors, however, the posterior median value for transmission 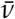 is very different from the experimental median. For the model with experiment-based priors, the posterior median is roughly an order of magnitude higher than the experimental median, and there is no overlap in the 95% credible intervals on 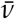 in the two cases. For the model with all vague priors, the posterior median is almost two orders of magnitude higher than the experimental median, and there is no overlap in the 95% credible interval on 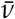 for that model with the 95% credible interval on 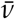 from the experimental data, or with the 95% credible interval on 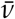 for the model with experiment-based priors (fig. 4). For models with experiment-based priors on parameters other than 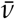, the posterior medians of 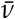 are similarly higher than the experimental median, or the posterior medians for any models with experiment-based priors on 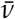 (fig. 5).

**Figure 5:**
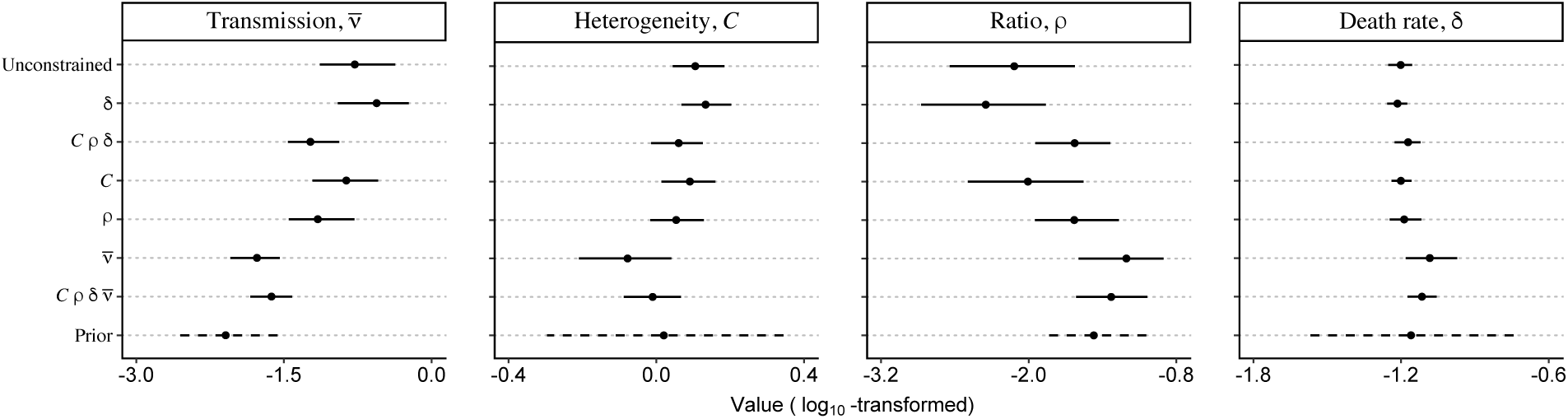
Summaries of the marginal posterior parameter estimates (columns) for all models (rows), compared to the experiment-based prior distributions derived from experiments (dashed lines). Dots are median values, while the horizontal bars show the 95% credible intervals. Note that each panel has a different scale on its horizontal axis.

For the models with experiment-based priors on transmission, the posterior distributions of transmission are thus concentrated at much lower values than the corresponding posteriors for the models with vague priors on transmission. This effect occurs because of the constraining effects of the experiment-based prior on transmission. The same models, however, are also strongly constrained by the epizootic data, and so their posterior distributions of transmission parameters are concentrated at much higher values than the prior itself. The posterior distributions of transmission rates for these models thus reflect the combined influence of the priors and the likelihood.

Part of the reason why models with experiment-based priors on transmission still fit the epizootic data reasonably well has to do with the effects of stochasticity on infection rates. As we show in the Online Appendix, high stochasticity can also increase overall transmission. Because the experiment-based priors on the transmission rate 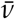 are centered at low transmission rates, models that use experiment-based priors on transmission attempt to explain the epizootic data using high stochasticity. The reasonably high *r*^2^ value for the model with experiment-based priors on most parameters is therefore due to high levels of transmission stochasticity. Although it may be that baculovirus epizootics in nature are indeed strongly affected by stochasticity, a more parsimonious explanation is that there is a mechanism operating in epizootics that is not in the model.

Comparison of WAIC scores then shows that the model with vague priors provides a much better explanation for the data than the models with experiment-based priors on the transmission rate (Table 3), with ΔWAIC *>* 5 in all cases. We therefore conclude that, in this pathogen, small-scale transmission is insufficient to explain large-scale epizootics.

**Table 3:**
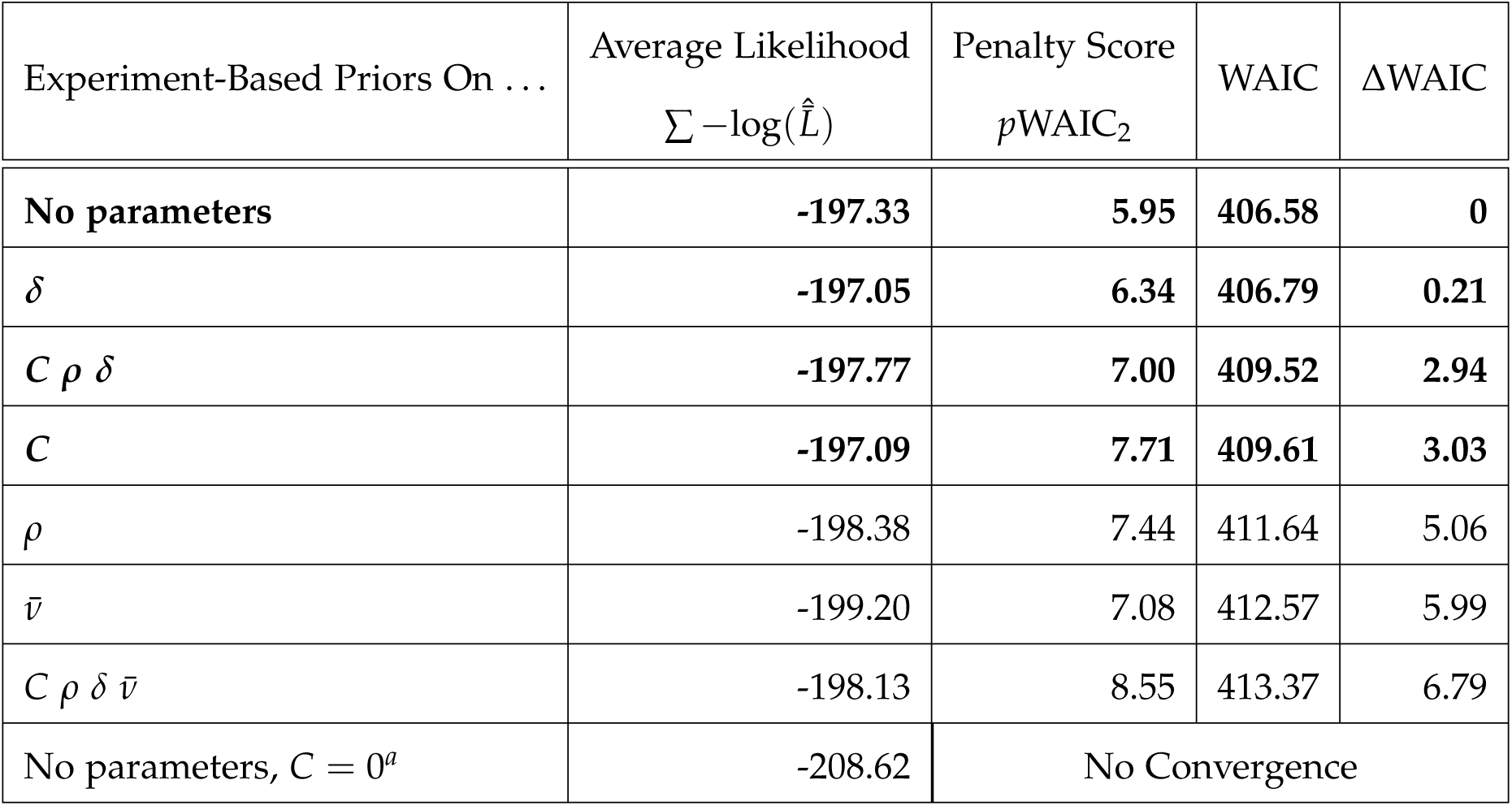
WAIC model selection for observational data. Models for which ΔWAIC*<≈* 3 are considered to be indistinguishable from the best model, and are therefore shown in bold face. *^a^*Because the model with no heterogeneity in transmission (*C* = 0) did not converge, the average likelihood for that model is a rough estimate based on non-converged MCMC samples.

| Experiment-Based Priors On ... | Average Likelihood<br>$\sum -\log(\hat{L})$ | Penalty Score<br>$p\text{WAIC}_2$ | WAIC | $\Delta\text{WAIC}$ |
| --- | --- | --- | --- | --- |
| **No parameters** | **-197.33** | **5.95** | **406.58** | **0** |
| $\delta$ | **-197.05** | **6.34** | **406.79** | **0.21** |
| $C \rho \delta$ | **-197.77** | **7.00** | **409.52** | **2.94** |
| $C$ | **-197.09** | **7.71** | **409.61** | **3.03** |
| $\rho$ | -198.38 | 7.44 | 411.64 | 5.06 |
| $\bar{v}$ | -199.20 | 7.08 | 412.57 | 5.99 |
| $C \rho \delta \bar{v}$ | -198.13 | 8.55 | 413.37 | 6.79 |
| No parameters, $C = 0^a$ | -208.62 | No Convergence | | |

That is not to say, however, that individual-level mechanisms do not play a role in epizootics. Evidence in support of the role of individual-level mechanisms comes from models with experiment-based priors on parameters other than transmission. For these models, ΔWAIC scores were less than 3, indicating that the fit of these models is effectively indistinguishable from the fit of the model with all vague priors. In the Online Appendices, we show that the visual fit of these models to the data is very similar to the visual fit of the best model, for which all priors were vague.

Of particular note is that the posterior medians for the model with all vague priors, and for the models with experiment-based priors on heterogeneity but not on transmission, are close to the posterior median for heterogeneity from our experimental data. We therefore conclude that individual heterogeneity in transmission plays a key role in the dynamics of the baculovirus. Overall, then, our results show that processes beyond the branch-scale affect epizootics, but that branch-scale processes also play an important role.

An important feature of host-pathogen models with high heterogeneity is that they predict lower infection rates at high host density, due to the dominating effects of resistant individuals, and higher infection rates at low host density, due to the presence of at least a few highly susceptible individuals (Dwyer et al., 1997). In figs. 2 and 3, infection rates were high across a broad range of densities in both the data and the models, consistent with these effects. Also because of these effects, models that do not account for heterogeneity provide poor fits to data from populations at either very low or very high densities (Table 3, also see Online Appendices).

Taken together, these results provide a complicated answer to our original question: are interactions between individual hosts on single branches sufficient to explain baculovirus epizootics in entire forests? The large differences in posterior values of transmission 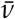 between the model with all vague priors, and the models with experiment-based priors on transmission, as well as the worse WAIC scores of models with experiment-based priors on transmission, suggest that there are processes affecting epizootics besides interactions between hosts on single branches, and thus that the answer to our question is no. The ability of models with experiment-based priors on heterogeneity in transmission nevertheless emphasizes that branch-scale processes also play a key role.

## Discussion

The assumption that interactions between individual hosts at a small scale determine infection and parasitization rates has been fundamental to studies of host-pathogen and host-parasitoid interactions for decades (Anderson and May, 1979; Varley et al., 1973). A common approach to understanding pathogen or parasite dynamics is therefore to estimate transmission rates from small-scale data or laboratory data (Blackwood et al., 2013; Buhnerkempe et al., 2011; George et al., 2011). Our results for the baculovirus of the Douglas-fir tussock moth instead show that estimating transmission from small-scale data provides a meaningfully worse fit to large-scale data than if transmission was estimated from the large-scale data alone. Our work therefore suggests that small-scale interactions between hosts are insufficient to explain the dynamics of this pathogen.

Direct tests of general models require specific biological systems, but we nevertheless argue that our results are of general significance. The basis of our argument is that, among animals, environmentally transmitted pathogens may be the rule rather than the exception (Cory and Myers, 2003; Duffy and Sivars-Becker, 2007; Mihaljevic et al., 2018; Rohani et al., 2003). Our results then suggest that, for such diseases, models that include only small-scale interactions between hosts may often be insufficient.

Our work does not definitively identify spatial structure as the missing mechanism in our models, but the failure of models that rely on branch-scale estimates of transmission at least suggests that the missing processes in our models operate at larger scales than the scale of our experiment. Moreover, there are several important factors that likely affect the pathogen but that are not included in our models, and each of these factors involves spatial structure or environmental heterogeneity.

First, Douglas-fir tussock moth larvae can grow and develop on multiple different host tree species, and forest tree-species composition varies strongly across the insect’s range (Shepherd et al., 1988). In British Columbia, Douglas fir is the dominant host tree species, while *Abies* species predominate in California and Nevada, with intermediate frequencies in other parts of the western USA. This is important because previous work showed that the transmission of the gypsy moth baculovirus can be strongly affected by variation in plant foliage chemistry (Elderd et al., 2013). If similar effects occur in the tussock moth baculovirus, differences in forest tree-species composition may have modulated epizootics in a way that was not accounted for in the model with experiment-based priors.

Second, although all 3 of the baculovirus isolates in our experiments were of the multicapsid or *Op*MNPV morphotype, in which viral capsids occur in clumps within occlusion bodies, there is a second, unicapsid or *Op*SNPV morphotype that occurs in tussock moth populations in nature, in which viral capsids occur singly within occlusion bodies (Hughes and Addison, 1970). The frequencies of the two morphotypes appear to vary latitudinally, with high frequencies of *Op*MNPV in British Columbia, high frequencies of *Op*SNPV in New Mexico, and intermediate frequencies in Washington, Oregon, Idaho, and California (Williams et al., 2011). Although not much is known about differences in phenotypes between the morphotypes, phylogenetic analyses have shown that the two are at least moderately diverged (Jakubowska et al., 2007), and it therefore seems likely that the phenotypes of the two morphotypes differ. This seems especially likely given that we observed meaningful differences in transmission parameters even within the three *Op*MNPV strains that we used in our experiment. Variation in morphotype frequency is thus a second possible missing mechanism in the model with experiment-based priors, while interactions between morphotypes and host-tree species provide yet a third possible missing mechanism.

Finally, tussock moth larvae are often observed to be at higher densities near the tops of trees, and this aggregation may increase infection rates relative to our experiments (Dwyer and Elkinton, 1993). Superficially, it seems unlikely that this mechanism plays a key role, because our estimates of over-dispersion levels are modest, but theory has shown that modest clumping can sometimes have strong effects (Bolker and Pacala, 1999). Clumping is therefore a final possible missing mechanism in the model with experiment-based priors.

Likely explanations for the missing mechanisms in the model thus have largely to do with spatial structure. We therefore advocate the further development of spatial theory in disease ecology. In particular, spatial models in disease ecology have often focused on traveling waves and other dramatic spatial phenomena (Dwyer, 1992), reflecting the focus of spatial models in ecology as a whole (Murray, 1989). Our work in contrast suggests that an unresolved question is, how do spatial patchiness and environmental heterogeneity together drive temporal dynamics? This is a long-standing problem in ecology (Bolker and Pacala, 1999), but our work suggests that solutions to the problem may have practical applications in pest control.

There are also two ways in which our work emphasizes the importance of stochasticity in pathogen dynamics. First, all of our models invoke substantial stochasticity to fit the epizootic data. The models with experiment-based priors on transmission have particularly high posterior estimates of stochasticity, not only because randomness helps those models better fit the epizootic data, but also because higher stochasticity by itself increases infection rates (in the Online Appendices, we prove this assertion). This effect occurs because increased stochasticity in transmission increases the frequency of both very low and very high transmission rates, but higher transmission rates have disproportionately stronger effects on the infection rate. For the models with experiment-based priors on transmission, the fitting routine therefore attempted to fit the epizootic data using high levels of stochasticity. This leads to more uncertain predictions, which is part of the reason why those models have larger (worse) WAIC scores.

Second, it is not clear that our models include the correct type of stochasticity. To explain this, we note that, because we fit a separate value of stochasticity to each population, it is possible to consider how stochasticity varied with host density. In the unsprayed populations in particular, the median posterior values of stochasticity were smaller in populations with higher initial host densities (Online Appendix). The stochasticity associated with small population sizes, known as “demographic stochasticity” (Bolker, 2008), may therefore have been more important than the environmental stochasticity that we included in our models. Because similar effects did not occur in the sprayed populations, we suspect that any such demographic stochasticity has do to with low initial densities of the pathogen, rather with low initial densities of hosts. Moreover, it seems likely that any such demographic stochasticity is compounded by the effects of space, because the number of occlusion bodies on a branch is of course much smaller than the total number of occlusion bodies in a forest.

In making these points, we are not arguing that a lack of consideration of demographic stochasticity means that our results were flawed, because we suspect that allowing for demographic stochasticity instead of environmental stochasticity would have given similar results. Our larger point is instead that further development of spatial models should also include careful consideration of the effects of stochasticity, and how stochasticity is compounded by spatial structure.

Although branch-scale transmission is insufficient to explain the dynamics of the Douglas-fir tussock moth baculovirus, it is important to remember that models with experiment-based priors on heterogeneity in transmission fit the data nearly as well as the model with vague priors. Individual-level mechanisms thus also play a key role in the dynamics of this pathogen. In disease ecology, host heterogeneity is typically only invoked in studies of sexually transmitted infections of humans (Keeling and Rohani, 2008), but our work suggests that host variation may have effects in many systems. Detecting such effects, however, may require a consideration of individual-scale data, as emphasized by Murdoch et al. (2005). Although individual data are unavailable in many host-pathogen systems, recent work has used measurements of antibody kinetics in individual hosts to estimate the force of infection (Pepin et al., 2017). A similar approach may allow for estimation of heterogeneity in host transmission.

Our estimates of heterogeneity *C* are also relevant to insect-pathogen population cycles. As fig. 4 shows, a substantial fraction of our posterior estimates of *C* are greater than 1, and for the model with vague priors, the lower bound on the 95% credible interval is above 1. This is important because, in simple, long-term models of insect outbreak cycles, values of heterogeneity *C >* 1 guarantee a stable point equilibrium (Dwyer et al., 2000). In such models, however, *C >* 1 can instead allow cycles if resistance is heritable, so that selection by the virus drives fluctuations in resistance (Elderd et al., 2008). Given that there is overwhelming evidence that Douglas-fir tussock moth populations have cyclic outbreaks (Mason, 1996), our estimates of heterogeneity suggest that selection plays a role in tussock moth population cycles, much as selection plays a role in gypsy moth population cycles (Páez et al., 2017).

Our Bayesian approach allowed us to show that our small-scale experimental data are not sufficient to explain the dynamics of the tussock moth baculovirus at large scales, even though the model with experiment-based priors fits the data fairly well. We therefore echo Restif et al. (2012)’s argument that Bayesian methods can allow for deep insights into disease dynamics. Moreover, in ecology, mechanistic model-fitting and high-performance computing are typically applied only to observational data (Ionides et al., 2015). This is problematic partly because a reliance on observational data alone can lead to flawed inferences (Cobey and Baskerville, 2016), but more broadly because mechanistic model-fitting is rarely used in experimental field ecology. By using model-fitting and high-performance computing to synthesize experimental and observational data, we hope to have shown that such tools can indeed be useful in experimental field ecology. The computational methods that we present here may therefore be of general usefulness.

## Acknowledgments

We are grateful for the support of dedicated and talented field technicians: Ruby An, Kate Lynne Logan, Rachel Hosman, Hannah Koch, Jenni Novak, Katherine Sirianni, Jeffrey Thorburn, Alison Hunter, Cara Skalisky, Alyssa Taylor, Jason Sims, and Rita Bennett. Roy Magelssen provided information on spray projects in general and some field support for the 2010 project on the Methow Ranger District. Epizootic data were collected with the assistance of Tom Eckberg, Idaho Department of Lands and Rebecca Powell, Forest Health Protection, Rocky Mountain Region. JRM was funded by a US Department of Agriculture (USDA) National Institute of Food and Agriculture (NIFA) Postdoctoral Fellowship (2014-67012-22272). Additional work was funded by the Okanogan-Wenatchee National Forest (Methow Ranger District), and a grant to KMP and GD from the USDA Forest Service Pesticide Impacts Assessment Program. Helpful feedback was provided by Iral Ragenovich, USDA Forest Service, Region 6 Forest Health Protection. Imre Otvos kindly provided the epizootic data from 1987. M.E. Martignoni provided important encouragement to G.D. at a crucial career juncture decades ago.

## Literature Cited

Anderson, R. M., and R. M. May. 1979. Population biology of infectious diseases: Part i. Nature 280:361–367.

Anderson, R. M., and R. M. May. 1980. Infectious-diseases and population-cycles of forest insects. Science 210:658–661.

Anderson, R. M., and R. M. May. 1992. Infectious diseases of humans: dynamics and control. Oxford University Press, Oxford.

Blackwood, J. C., D. G. Streicker, S. Altizer, and P. Rohani. 2013. Resolving the roles of immunity, pathogenesis, and immigration for rabies persistence in vampire bats. Proceedings of the National Academy of Sciences 110:20837–20842.

Bolker, B. M. 2008. Ecological models and data in R. Princeton University Press.

Bolker, B. M., and S. W. Pacala. 1999. Spatial moment equations for plant competition: understanding spatial strategies and the advantages of short dispersal. The American Naturalist 153:575–602.

Buhnerkempe, M. G., R. J. Eisen, B. Goodell, K. L. Gage, M. F. Antolin, and C. T. Webb. 2011. Transmission shifts underlie variability in population responses to yersinia pestis infection. PloS one 6:e22498.

Burand, J. P., and E. J. Park. 1992. Effect of nuclear polyhedrosis-virus infection on the development and pupation of gypsy-moth larvae. Journal of Invertebrate Pathology 60:171–175.

Cobey, S., and E. B. Baskerville. 2016. Limits to causal inference with state-space reconstruction for infectious disease. PloS one 11:e0169050.

Cory, J. S., and K. Hoover. 2006. Plant-mediated effects in insect-pathogen interactions. Trends in Ecology and Evolution 21:278–286.

Cory, J. S., and J. H. Myers. 2003. The ecology and evolution of insect baculoviruses. Annual Reviews of Ecology and Systematics 34:239–272.

Cox, D. R., and E. Snell. 1989. Analysis of binary data. Routledge.

Duffy, M. A., and L. Sivars-Becker. 2007. Rapid evolution and ecological host-parasite dynamics. Ecology Letters 10:44–53.

Dwyer, G. 1991. The effects of density, stage and spatial heterogeneity on the transmission of an insect virus. Ecology 72:559–574.

Dwyer, G. 1992. On the spatial spread of insect pathogens - theory and experiment. Ecology 73:479– 494.

Dwyer, G., J. Dushoff, J. S. Elkinton, and S. A. Levin. 2000. Pathogen-driven outbreaks in forest defoliators revisited: Building models from experimental data. American Naturalist 156:105– 120.

Dwyer, G., and J. S. Elkinton. 1993. Using simple-models to predict virus epizootics in gypsymoth populations. Journal Of Animal Ecology 62:1–11.

Dwyer, G. 1995. Host dispersal and the spatial spread of insect pathogens. Ecology 76:1262–1275.

Dwyer, G., J. S. Elkinton, and J. P. Buonaccorsi. 1997. Host heterogeneity in susceptibility and disease dynamics: Tests of a mathematical model. American Naturalist 150:685–707.

Eakin, L., M. Wang, and G. Dwyer. 2015. The effects of the avoidance of infectious hosts on infection risk in an insect-pathogen interaction. Am. Nat. 185:pp. 100–112.

Elderd, B., V. Dukic, and G. Dwyer. 2006. Uncertainty in predictions of disease spread and public-health responses to bioterrorism and emerging diseases. Proceedings of the National Academy of Sciences 103:15693–15697.

Elderd, B. D. 2013. Developing models of disease transmission: Insights from ecological studies of insects and their baculoviruses. PLoS Pathogens 9.

Elderd, B. D., J. Dushoff, and G. Dwyer. 2008. Host-pathogen interactions, insect outbreaks, and natural selection for disease resistance. The American Naturalist 172:829–842.

Elderd, B. D., B. J. Rehill, K. J. Haynes, and G. Dwyer. 2013. Induced plant defenses, host– pathogen interactions, and forest insect outbreaks. Proc. Natl. Acad. Sci. 110:14978–14983.

Fleming-Davies, A. E., V. Dukic, V. Andreasen, and G. Dwyer. 2015. Effects of host heterogeneity on pathogen diversity and evolution. Ecology Letters 18:1252–1261.

Fuller, E., B. D. Elderd, and G. Dwyer. 2012. Pathogen persistence in the environment and insect-baculovirus interactions: Disease-density thresholds, epidemic burnout, and insect outbreaks. Am. Nat. 179:pp. E70–E96.

Gelman, A., J. B. Carlin, H. S. Stern, D. B. Dunson, A. Vehtari, and D. B. Rubin. 2014. Bayesian Data Analysis, Third Edition. Chapman & Hall/CRC Press. New York, NY.

George, D. B., C. T. Webb, M. L. Farnsworth, T. J. O’Shea, R. A. Bowen, D. L. Smith, T. R. Stanley, L. E. Ellison, and C. E. Rupprecht. 2011. Host and viral ecology determine bat rabies seasonality and maintenance. Proceedings of the National Academy of Sciences 108:10208–10213.

Grove, M. J., and K. Hoover. 2007. Intrastadial developmental resistance of third instar gypsy moths (*Lymantria dispar* l.) to L. dispar nucleopolyhedrovirus. Biological Control 40:355–361.

Hall, S. R., J. L. Simonis, R. M. Nisbet, A. J. Tessier, and C. E. Cáceres. 2009. Resource ecology of virulence in a planktonic host-parasite system: an explanation using dynamic energy budgets. The American Naturalist 174:149–162.

Hamede, R. K., J. Bashford, H. McCallum, and M. Jones. 2009. Contact networks in a wild tasmanian devil (sarcophilus harrisii) population: using social network analysis to reveal seasonal variability in social behaviour and its implications for transmission of devil facial tumour disease. Ecology letters 12:1147–1157.

Han, X., and P. E. Kloeden. 2017. Random Ordinary Differential Equations and Their Numerical Solution. Springer.

Hughes, K., and R. Addison. 1970. Two nuclear polyhedrosis viruses of Douglas-fir tussock moth. Journal of Invertebrate Pathology 16:196–&.

Hunter-Fujita, F. R., P. F. Entwistle, H. F. Evans, and N. E. Crook. 1998. Insect viruses and pest management. John Wiley and Sons: Somerset, New Jersey.

Ionides, E. L., D. Nguyen, Y. Atchadé, S. Stoev, and A. A. King. 2015. Inference for dynamic and latent variable models via iterated, perturbed Bayes maps. Proceedings of the National Academy of Sciences 112:719–724.

Jakubowska, A., M. M. van Oers, I. S. Otvos, and J. M. Vlak. 2007. Phylogenetic analysis of orgyia pseudotsugata single-nucleocapsid nucleopolyhedrovirus. Virologica Sinica 22:257–265.

Keeling, M. J., and P. Rohani. 2008. Modeling Infectious Diseases in Humans and Animals. Princeton University Press.

Kermack, W., and A. McKendrick. 1927. A contribution to the mathematical theory of epidemics. Proceedings of the Royal Society of London, Series A 115:700–721.

King, A. A., E. L. Ionides, M. Pascual, and M. J. Bouma. 2008. Inapparent infections and cholera dynamics. Nature 454:877–U29.

Kuznetsova, A., D. McKenzie, P. Banser, T. Siddique, and J. M. Aiken. 2014. Potential role of soil properties in the spread of cwd in western canada. Prion 8:92–99.

Martignoni, M. E. 1999. History of tm biocontrol-1, the first registered virus-based produced for control of a forest insect. The American Entomologist 45:30–37.

Mason, R. 1996. Dynamic behavior of Douglas-fir tussock moth populations in the Pacific north-west. Forest Science 42:182–191.

Mason, R., and T. Torgersen. 1983. Mortality of larvae in stocked cohorts of the Douglas-fir tussock moth, *Orgyia pseudotsugata* (Lepidoptera: Lymantriidae). Canadian Entomologist 115:1119–1127.

McCallum, H. 2016. Models for managing wildlife disease. Parasitology 143:805–820.

McCullagh, P., and J. Nelder. 1989. Generalized Linear Models. Chapman & Hall, Boca Raton, FL.

Mihaljevic, J. R., J. T. Hoverman, and P. T. Johnson. 2018. Co-exposure to multiple ranavirus types enhances viral infectivity and replication in a larval amphibian system. Diseases of aquatic organisms 132:23–35.

Miller, L. K. 1997. The baculoviruses. Plenum Press.

Moreau, G., and C. J. Lucarotti. 2007. A brief review of the past use of baculoviruses for the management of eruptive forest defoliators and recent developments on a sawfly virus in canada. Forestry Chronicle 83:105–112.

Murdoch, W., C. J. Briggs, and S. Swarbrick. 2005. Host suppression and stability in a parasitoid-host system: experimental demonstration. Science 309:610–613.

Murray, J. D. 1989. Mathematical biology, vol. 19 of biomathematics.

Øksendal, B. 2003. Stochastic differential equations. Pages 65–84 *in* Stochastic differential equations. Springer.

Otvos, I., J. Cunningham, and R. Alfaro. 1987. Aerial Application Of Nuclear Polyhedrosis-Virus Against Douglas-Fir Tussock Moth, Orgyia-Pseudotsugata (Mcdunnough) (Lepidoptera, Lymantriidae).2. Impact 1-Year And 2 Years After Application. Canadian Entomologist 119:707–715.

Otvos, I. S., J. C. Cunningham, and L. M. Friskie. 1987. Aerial application of nuclear polyhedrosis virus against douglas-fir tussock moth, Orgyia pseudostugata (Mcdunnough) (Lepidoptera: Lymantriidae). 1. impact in the year of application. Canadian Entomologist 119:697–706.

Páez, D., V. Dukic, J. Dushoff, A. Fleming-Davies, and G. Dwyer. 2017. Effects of pathogen exposure on life-history variation in the gypsy moth (lymantria dispar). The American Naturalist accepted pending minor revision.

Páez, D., A. Fleming-Davies, and G. Dwyer. 2015. Effects of pathogen exposure on life-history variation in the gypsy moth (lymantria dispar). Journal of evolutionary biology 28:1828–1839.

Parker, B. J., B. D. Elderd, and G. Dwyer. 2010. Host behaviour and exposure risk in an insect-pathogen interaction. Journal of Animal Ecology 79:863–870.

Pascual, M. A., and P. Kareiva. 1996. Predicting the outcome of competition using experimental data: maximum likelihood and bayesian approaches. Ecology 77:337–349.

Pepin, K. M., S. L. Kay, B. D. Golas, S. S. Shriner, A. T. Gilbert, R. S. Miller, A. L. Graham, S. Riley, P. C. Cross, M. D. Samuel, et al. 2017. Inferring infection hazard in wildlife populations by linking data across individual and population scales. Ecology Letters 20:275–292.

Polivka, K., G. Dwyer, and C. Mehmel. 2017. Environmental persistence of a pathogen used in microbial insect control. Research Note PNW-RN-573, Pacific Northwest Research Station, USDA Forest Service.

Press, W. H., S. A. Teukolsky, W. T. Vetterling, and B. P. Flannery. 1992. Numerical recipes in C, vol. 2. Cambridge university press Cambridge.

Restif, O., D. T. Hayman, J. R. Pulliam, R. K. Plowright, D. B. George, A. D. Luis, A. A. Cunningham, R. A. Bowen, A. R. Fooks, T. J. O’Shea, et al. 2012. Model-guided fieldwork: practical guidelines for multidisciplinary research on wildlife ecological and epidemiological dynamics. Ecology letters 15:1083–1094.

Rohani, P., C. J. Green, N. B. Mantilla, and B. T. Grenfell. 2003. Ecological interference between fatal diseases. Nature 422:885–888.

Rohrmann, G. F. 2014. Baculovirus nucleocapsid aggregation (MNPV vs SNPV): an evolutionary strategy, or a product of replication conditions? Virus Genes 49:351–357.

Ross, S. 2002. Simulation, 3rd. Edition. Academic Press, New York.

Schindelin, J., C. T. Rueden, M. C. Hiner, and K. W. Eliceiri. 2015. The ImageJ ecosystem: An open platform for biomedical image analysis. Molecular Reproduction and Development 82:518–529.

Shepherd, R., D. Bennett, J. Dale, S. Tunnock, R. Dolph, and R. Thier. 1988. Evidence of synchronized cycles in outbreak patterns of Douglas-fir tussock moth, *Orgyia pseudotsugata* (McDunnough) (Lepidoptera:Lymantriidae). Memoirs of the Entomological Society of Canada pages 107–121.

Shepherd, R. F., I. S. Otvos, R. J. Chorney, and J. C. Cunningham. 1984. Pest-management of Douglas-fir tussock moth (Lepidoptera: Lymantriidae) - prevention of an outbreak through early treatment with a nuclear polyhedrosis-virus by ground and aerial applications. Canadian Entomologist 116:1533–1542.

Shocket, M. S., A. T. Strauss, J. L. Hite, M. Šljivar, D. J. Civitello, M. A. Duffy, C. E. Cáceres, and S. R. Hall. 2018. Temperature drives epidemics in a zooplankton-fungus disease system: A trait-driven approach points to transmission via host foraging. The American Naturalist 191:435–451.

Somerville, R. A., K. Fernie, A. Smith, K. Bishop, B. C. Maddison, K. C. Gough, and N. Hunter. 2019. Bse infectivity survives burial for five years with only limited spread. Archives of virology 164:1135–1145.

Thompson, C., and D. Scott. 1979. Production And Persistence Of The Nuclear Polyhedrosis-Virus Of The Douglas-Fir Tussock Moth, Orgyia-Pseudotsugata (Lepidoptera, Lymantriidae), In The Forest Ecosystem. Journal Of Invertebrate Pathology 33:57–65.

Varley, G. C., G. R. Gradwell, and M. P. Hassell. 1973. Insect population ecology: an analytical approach. Blackwell Scientific Publications: Oxford.

Williams, H. L., K. S. Monge-Monge, I. S. Otvos, R. Reardon, and I. Ragenovich. 2011. Genotypic variation among Douglas-fir tussock moth nucleopolyhedrovirus (OpNPV) isolates in the western United States. Journal of Invertebrate Pathology 108:13–21.

