## Supplementary Materials for "An empirical test of the role of small-scale transmission in large-scale disease dynamics"

Mihaljevic *et al.*

### Contents

**1 Epizootic data** **1**

**2 Speed of kill experiment** **4**

**3 Experimental estimate of the ratio parameter, $\rho$** **6**

**4 Experiment-based Priors** **8**

**5 Statistical Complications in our Estimation of Average Transmission** **8**

**6 Effects of stochasticity in the model** **9**

**7 Numerical Algorithms and Model Selection** **12**

**8 Visual fits to the observational data for models with intermediate numbers of experiment-based priors.** **18**

**9 Marginal posteriors all models** **23**

**10 JAGS model statement for fitting the transmission model to experimental data** **32**

**11 JAGS model statement to fit a Gamma distribution to the time-to-death data** **35**

### 1 Epizootic data

#### *Epizootics*

The number of larvae collected varied between studies due to differences in overall design, initial host density, and the resources available for collections. Also, near the end of epizootics, sample sizes usually dropped because high baculovirus mortality made it hard to find live larvae (cadavers disintegrate too rapidly to allow for meaningful collections and cadaver collection would create a high risk of cross contamination among sampled larvae). The number of larvae collected per week therefore ranged from 7 to 197, with a mean of 95.6, and with a total across sites of 5830 (Otvos et al., 1987; Polivka et al., 2012; Scott and Spiegel, 2002). In fitting our models, we then had 61 observations of virus infection rates, 30 at control sites and 31 at sprayed sites. Initial host densities spanned almost two orders of magnitude, ranging from 7.20 to 232.0 larvae/$m^2$, with a mean of 103.2. The control and sprayed sites had similar ranges and means of initial larval host density (table A1).

In two cases, one the work of Otvos et al. (1987) in the Kamloops Forest District of British Columbia, and one the work of a co-author (C.J.M.) in the Methow Valley of Washington State, data were collected both from sprayed treatment plots (Otvos et al.: 4 sites; C.J.M.: 1 site), and control sites (Otvos et al.: 3 sites; C.J.M.: 2 sites, reported in Polivka et al. 2012). One of us (K.M.P.) also collected data from two naturally occurring epizootics, one in the Lovell Valley (Benewah Co.), Idaho, and one in Cheyenne Mountain State Park in Colorado (table A1).

Initial host density in our observation sites was measured using one of three methods, depending on the site. The first two methods consisted of standardized cocoon or egg mass surveys in the fall or spring preceding the insect collection, respectively (Mason et al., 1993). Cocoon surveys were conducted for the 2010 Washington sites, while an egg mass survey was conducted for the Idaho site. In these surveys, 25 trees were randomly selected across a 0.8 to 2.0 *ha* area, and the number of cocoons or egg masses on the underside of three 45 *cm* branch tips in the lower

| Site | Location | Year | Initial Host Density (Larvae $\cdot$ $m^{-2}$) |
|---|---|---|---|
| C1 | Washington | 2010 | 7.20 |
| C2 | British Columbia | 1987 | 44.35 |
| C3 | Colorado[a] | 2015 | 47.85 |
| C4 | British Columbia | 1987 | 112.09 |
| C5 | Idaho | 2010 | 126.08 |
| C6 | Washington | 2010 | 149.00 |
| C7 | British Columbia | 1987 | 165.39 |
| T1 | Washington | 2010 | 8.99 |
| T2 | British Columbia | 1987 | 34.27 |
| T3 | British Columbia | 1987 | 124.08 |
| T4 | British Columbia | 1987 | 172.65 |
| T5 | British Columbia | 1987 | 232.03 |

Table A1: Labels and characteristics of sites used for the observational data set. C1, C2, and so on indicate unsprayed control sites, while T1, T2 and so on indicate sprayed sites. [a]The initial host density for the Colorado site was estimated from the data.

crown of each tree was counted (Mason et al., 1993). Because the pupae in the cocoons give rise to adult moths, we inferred the density of larvae resulting from cocoons by multiplying cocoon density by 0.5 to approximate the density of emerging females, and we inferred the initial larval density by multiplying by 145, the average number of larvae that hatch per egg mass (Mason et al., 1977). For the egg-mass survey data, we used the same calculation, except that of course we did not need to multiply by 0.5 to account for the fraction of pupae that produce females.

The third method consisted of larval surveys in the early spring preceding virus treatment, and was used by Otvos et al. (1987). In these surveys, 90 45 *cm* branch samples were taken from 45 trees at each site. The foliage area was estimated for each branch, and larvae were counted to yield larval density per $m^2$, which was then averaged across branches (Otvos et al., 1987). We did not have an estimate of initial host density for the 2015 control site in Colorado, and so for that site, we estimated the initial larval density along with the epidemiological parameters.

In some control sites, several weeks passed before sampling began, or before any infections were detected. In these cases, we adjusted the initial host density to account for non-disease mortality, due for example to predation or dessication, before the epizootic began. To simplify the problem, we assumed that this mortality rate was constant, and we estimated it using data from Mason and Torgersen (1983) on larval survivorship in two tussock moth populations. We fit a simple model of density-independent mortality to these data, using a binomial likelihood, which gives a log-likelihood score of:

$$-\sum_t (N(0) - N(t)) \log\left(1 - \mathrm{e}^{-bt}\right) + N(t) \log(\mathrm{e}^{-bt}). \tag{A1}$$

Here, the initial population size $N(0) = 150$, while $N(t)$ is the population size at $t$ days post hatching. To estimate the mortality rate $b$, we used the one-dimensional nonlinear fitting function `optimize()` in the `R` statistical language. The best-fit mortality rate is $b = 0.037$. As fig. A1 shows, the model provides a reasonable fit to the data, suggesting that the assumption of constant mortality is reasonable. Also, the model provided a good fit to data from both branches on which emigration was allowed, and on branches from which emigration was prevented, suggesting that mortality was mostly due to predation rather than to emigration. We used this estimate of $b$ to adjust the initial susceptible host density in epizootics for which the virus infection rate was zero for 2 or more weeks early in the larval period.

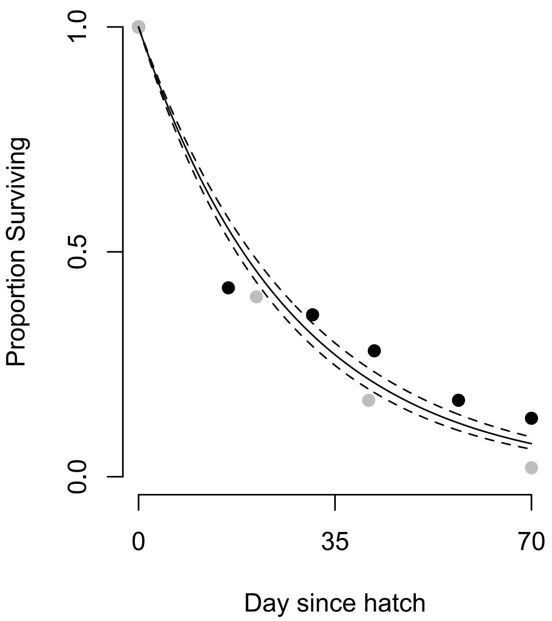

Figure A1: Fit of a density-independent mortality model to data from Mason and Torgersen (1983)'s fig 2. The black dots are data from experimental populations that were exposed to natural mortality factors, including predators, and for which emigration was allowed. Gray dots are data from adjacent populations for which emigration was prevented using cages that allowed access by small predators. The solid lines represent the maximum likelihood estimate of the decay function, while the dashed lines represent bootstrapped 95% confidence bounds.

#### 2 Speed of kill experiment

DFTM egg masses were collected from sites C1 and T1 (table A1), sterilized, and reared to the fourth instar following the same protocols as the uninfected larvae in the field transmission experiment. Larvae were then exposed to the TMB-1 isolate by placing them on Douglas-fir seedlings in the greenhouse, at a temperature of 30 °C. To apply the virus to the foliage, we prepared a solution of TMB-1 (Lot no. 4 USDA Forest Service, Corvallis, Oregon) at a concentration of 0.018g $\cdot l^{-1}$, equal to the concentration used in large microbial control projects (R. Magelssen, pers. comm.). We then sprayed the foliage with 4.5 mL, a quantity sufficient to cover all the foliage on the small trees that we used.

We allowed larvae to feed on the foliage for 7 days and then we transferred them individually to 2 oz cups with artificial diet. Larvae were then incubated at 27 °C and monitored daily for mortality. Larvae that died before pupation were autopsied to confirm virus death.

In the SEIR model, hosts proceed through several exposed classes, leading to a gamma-distributed speed of kill (see main text). To estimate the death rate parameter $1/\delta$, we therefore used Bayesian methods to fit the following gamma distribution to the speed of kill data;

$$f(x;\alpha,\beta) = \frac{\beta^{\alpha} x^{\alpha-1} \mathrm{e}^{-\beta x}}{\Gamma(\alpha)} \tag{A2}$$

In terms of the parameters $\alpha$ and $\beta$, as well as the parameters $m$ and $\delta$ of the SEIR model, the mean of this distribution is: $\mathbf{E}[X] = \frac{\alpha}{\beta} = \frac{1}{\delta}$ and the variance is: $\mathrm{Var}[X] = \frac{\alpha}{\beta^2} = \frac{1}{m\delta^2}$. Thus, by estimating $\alpha$ and $\beta$ from our speed of kill data, we were able to estimate the death rate parameter $\delta$ in our epidemiological model.

We fit the gamma distribution in JAGS (http://mcmc-jags.sourceforge.net/) using flat priors for $\alpha$ and $\beta$, three MCMC chains, an adaptation period of 10,000 iterations, and 50,000 sampling iterations, thinning by 50, for a total of 1000 samples from the posterior. The resulting distribution explains most of the variation in the data (fig. A2). From the posterior distribution of the parameters, the mean $\alpha = 28.84$, with variance = 42.6, while the mean $\beta = 1.9$, with variance = 0.2. Given the relationship between these parameters and $\delta$, our posterior mean $\delta = 0.067$, so that the speed of kill is approximately 15 days.

By allowing larvae to feed freely on virus-contaminated foliage, we attempted to ensure that the doses that larvae consumed were similar to the doses that they consume in nature. A possible downside to this approach, however, is that larvae may have became infected over several days, whereas in laboratory experiments, larvae typically become infected over a few hours (Dwyer et al., 2005). Possibly for this reason, the variance in the speed of kill was substantially larger than

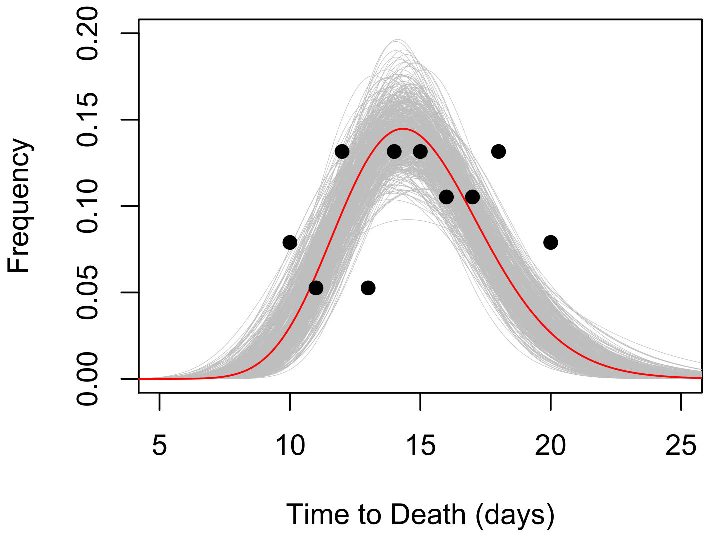

Figure A2: A comparison of the fitted gamma distribution to the speed of kill data. The black points represent the data, the red line represents the model's prediction based on the median values of the parameters, and the gray lines represent a set of model predictions using parameters drawn randomly from the joint posterior.

it was in a previous laboratory experiment (Dwyer, 1991), while nevertheless being extremely small compared to the mean (variance: $4.4 \times 10^{-6}$). The experimental data therefore imply a value of $m = 29$ (range 10-50), but because of the near-zero variance in previous work, in our fitting routine, we set $m = 200$. In fact, for values of $m$ much greater than 10, the variance in the speed of kill is very slight, so using $m = 29$ probably would have given similar results.

#### 3 Experimental estimate of the ratio parameter, $\rho$

To estimate the ratio parameter $\rho$, we infected 40 first instars and 20 fourth instars by exposing them to virus-contaminated diet in the lab, and then estimating the number of viral occlusion bodies (OBs) that were produced by each larva. We randomly assigned one of our three virus isolates to each larva, but to allow for variation across isolates, we pooled the data to produce an overall estimate of $\rho$. After exposure, larvae were reared individually at 26°C in 2*oz* cups until death, and then stored at -20°C in individual micro-centrifuge tubes. Thawed larvae were later vortexed in 5 mL of $dH_2O$ in the micro-centrifuge tubes, and then filtered through cheesecloth into a 50 mL centrifuge tube. After we added an additional 25 mL DI water to the tube, we centrifuged the solution for 10 minutes at 5000 $g$ to pellet the virus. The supernatant was then poured off and the viral pellet was re-suspended in 10 mL of DI water. We vortexed the solution for 3 minutes and counted OBs using a hemocytometer. We calculated an average density of OB/$\mu$L from two replicate counts of 10 $\mu$L aliquots of the virus isolate.

Viral densities ranged from 50 to 2150 OBs/$\mu$l for first instars, with a mean of 340, and from 1575 to 9775 OBs/$\mu$l for fourth instars, with a mean of 4832. Based on 1000 bootstraps of these data, we used the following log-normal prior distribution for $\rho$ in the field experiment: $\rho \sim LN(-2.77, 0.83)$.

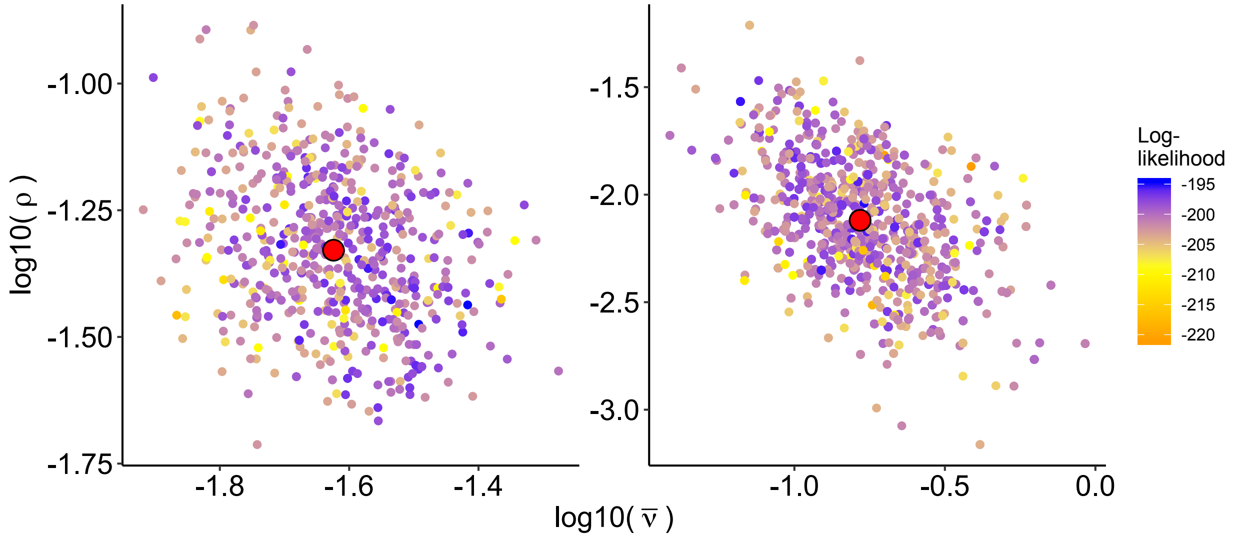

Figure A3: Joint marginal posterior of the ratio parameter, $\rho$, and average transmission rate, $\bar{\nu}$. The left panel is from the model with all experimentally-derived priors, while the right panel is from the model with all vague priors (unconstrained). The large red points represent the median marginal posterior values for each parameter.

We noticed that when we fit the model without the experimentally-based priors on $\rho$ or $\bar{\nu}$ (i.e., the unconstrained model), there seemed to be a pattern in which both parameter values shifted away from their experimentally-derived estimates, in a correlated way (see fig. 5 of the main text). To test if this was evidence of non-identifiability in either parameter in the case of vague priors, we plotted the parameters' joint marginal posterior. As can be seen, the correlations in the joint marginal posterior are relatively weak, showing no or very limited evidence of non-identifiability (fig. A3). Furthermore, there is no evidence of non-identifiability when both parameters are constrained by experimental data.

#### 4 Experiment-based Priors

The data from our field experiment gave the following prior probability distributions: $\bar{\nu} \sim LN(-4.82, 0.59)$, $k \sim LN(-0.094, 0.75)$, $\rho \sim LN(-3.39, 0.50)$, where $LN$ represents a log-normal distribution. For $k$ and $\rho$, we inflated the prior variance slightly to improve chain mixing, but the effects were minimal. When we measured the speed of kill of infected larvae in the laboratory, the average was $14.9d$, with very low variance (see above). The mean value of the death rate is then $\delta = 1/14.9 = 0.067$. Because our sample size was modest ($n = 38$ larvae), we inflated the variance on the prior for $\delta$; however, this precautionary variance inflation had little effect on our results.

#### 5 Statistical Complications in our Estimation of Average Transmission

First, as we mentioned, our failed efforts to estimate the decay rate $\mu$ experimentally suggest that $\mu$ may be very low for this virus (Dwyer, 1991), thus matching the posterior values of $\mu$ for the models with experiment-based priors on $\bar{\nu}$. Second, our experimental estimate of $\bar{\nu}$ depends on our estimate of the ratio parameter $\rho$, such that high values of $\rho$ lead to low estimates of $\bar{\nu}$. Estimating $\rho$ is difficult enough that such underestimation may have been a significant problem, and may explain why our experimental estimate of $\bar{\nu}$ appears to be inaccurately low. Third, the isolates present in the epizootic plots may have been phenotypically different from the isolates we used in our transmission experiment. Given the strong variation across isolates that is apparent in our transmission experiment, differences in isolates could lead to very different transmission rates between experiments and epizootics.

### 6 Effects of stochasticity in the model

In our model, the stochastic variation in transmission rates varied among sites, such that larger variation $\sigma$ increased the effective transmission rate over the course of the epizootic. Increased stochasticity thus led to more severe epizootics than expected from a deterministic model with the same parameters (fig. A4).

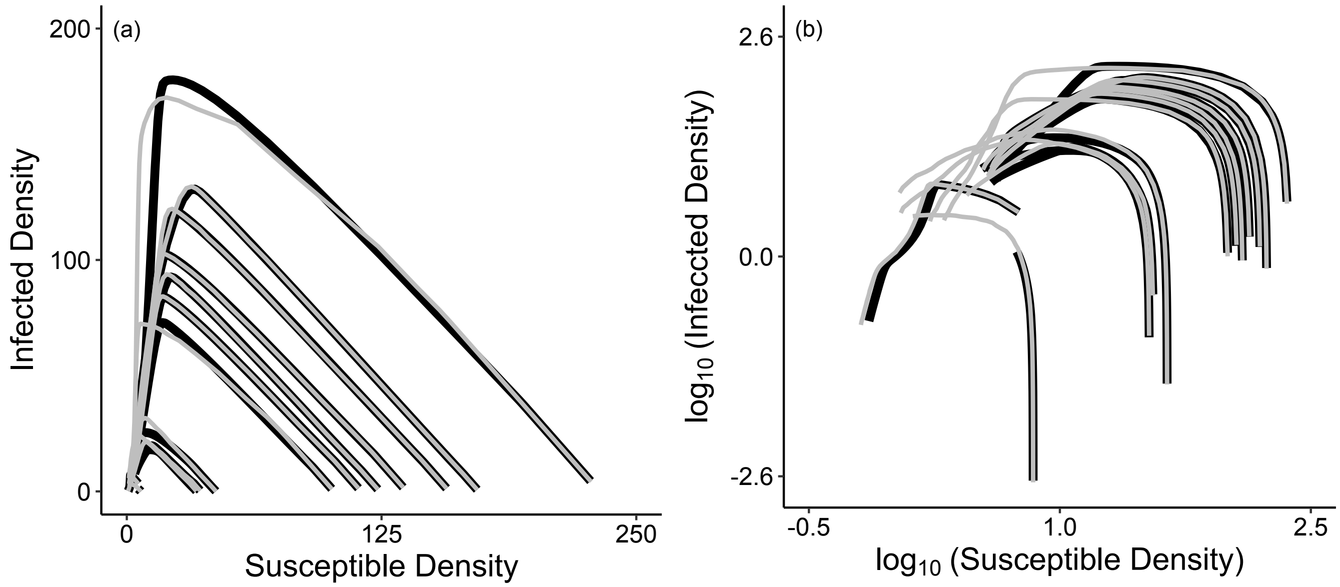

Figure A4: (a) Simulated phase portraits for our model, using initial conditions estimated from each of our twelve epizootics. The solid black lines represent trajectories from a deterministic version of the SEIR model, while the gray lines represent median trajectories over 500 realizations of the stochastic model, with time proceeding from right to left. The infected density in the model is measured as the sum of the densities in the exposed classes, $\sum E_m$. (b) Same as (a) but with log-transformed densities to illustrate more clearly that stochasticity leads to more severe epizootics.

To explain this effect, here we show that the mean transmission rate is always higher for the stochastic model. First, for the stochastic model, the average transmission rate $E[\bar{\nu}]$ can be expressed as:

$$\mathrm{E}\left[\nu_t\right] \quad = \mathrm{E}\left[\mathrm{e}^{\epsilon_t}\bar{\nu}\right] = \bar{\nu}\mathrm{E}\left[\mathrm{e}^{\epsilon_t}\right]. \tag{A3}$$

The rightmost expectation can be calculated according to,

$$\mathrm{E}\left[\mathrm{e}^{\epsilon_t}\right] \quad = \int \mathrm{e}^{\epsilon_t} d\mathrm{p}(\epsilon_t). \tag{A4}$$

In practice, we specify that the distribution of the $\epsilon_t$'s is normal, and so by Jensen's inequality we have;

$$\int_{-\infty}^{\infty} e^{\epsilon_t} dp(\epsilon_t) \quad = \int_{-\infty}^{\infty} e^{x} \frac{1}{\sqrt{2\pi\sigma^2}} e^{-\frac{x^2}{2\sigma^2}} dx > 1. \tag{A5}$$

We therefore have that,

$$E[\nu] \quad = \bar{\nu} E[e^{\epsilon_t}] > \bar{\nu}. \tag{A6}$$

In other words, the expected transmission rate of the stochastic model is always higher than the (constant) transmission rate of the deterministic model.

Interestingly, our estimates of the stochasticity term $\sigma$ vary in a consistent way with initial host density. As fig. A5 shows, the estimated values of $\sigma$ tended to be higher in plots with lower initial host densities, especially for control plots (fig. A5), and this trend is consistent across model types (see *Marginal Posterior* section, below, for parameter estimates across model types). This result suggests that what we are attributing to weather stochasticity may instead be stochasticity due to the chance events that befall individuals, known as "demographic stochasticity" (Bolker, 2008). In our case, transmission can be affected by behavioral decision-making and individual variation in exposure risk (Parker et al., 2010), either of which may be subject to chance at the individual level. Because demographic stochasticity has stronger effects in smaller populations, the higher values of $\sigma$ that we estimated in lower-density control plots may reflect demographic stochasticity. In sprayed plots in contrast, demographic stochasticity may have had weaker effects because the initial inundation of virus may have homogenized infection risk across space relative to the natural situation in which the virus is clumped into cadavers (D'Amico et al., 2005; Eakin et al., 2015). The spray may therefore have reduced or eliminated the effects of individual variation in behavior early in the epizootic.

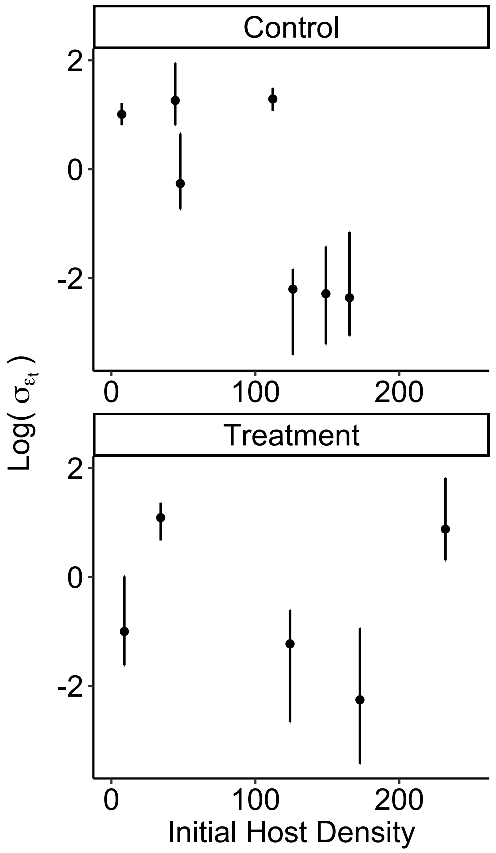

Figure A5: Relationship between stochastic transmission and initial host density for our 12 epizootics, separated into control and sprayed treatment plots. Error bars are 95% credible intervals.

### 7 Numerical Algorithms and Model Selection

Here we describe the numerical algorithms that we used to fit our SEIR model to the epizootic data, and the details of our model selection approach. First, as we mentioned in the main text, by using random ordinary differential equations instead of a stochastic ordinary differential equations, we were able to use a fairly simple numerical solution algorithm. Specifically, our algorithm solves the differential equations for each day in the epizootic, using a predictor-corrector algorithm. Specifically, we use a Runge-Kutta-Fehlberg algorithm that is a standard part of the Gnu Scientific Library. This algorithm compares 4th-order and 5th-order Runge-Kutta routines, and adjusts the time step to keep the difference between the two solutions reasonably low. Once this numerical routine is complete for one day, it re-sets the intial conditions to the ending conditions of that day, and then it calculates the solution for the next day. More sophisticated algorithms are available (Han and Kloeden, 2017), and such algorithms are probably faster. Our algorithm, however, had the advantage of being fairly obvious, which made implementing it straightforward. More importantly, it was fast enough for our purposes.

Given this numerical-solution algorithm, the fitting algorithm that we used is known as "line search-MCMC". Using line-search MCMC required that we carry out 3 steps, which can be briefly summarized as follows. First, we conducted a Bayesian line search through the multi-dimensional parameter space of the model, with multiple random re-starts of the line searches. Second, we performed principle components analysis (PCA) on the line-search results, in order to construct proposals for MCMC. We then used these proposals to implement an MCMC routine to sample from the posterior distribution of the model, and thus to fit the model to the epizootic data. The first and third steps required that we calculate model likelihood scores by averaging across the likelihoods of many model realizations, effectively integrating out the stochasticity across realizations. Here we explain these steps in more detail. All methods were executed using custom code in the C programming language (Kernighan and Ritchie, 2006).

#### 7.1 Line-search, Markov chain Monte Carlo (line-search MCMC)

The line-search MCMC method was first proposed by Kennedy et al. (2014) to adapt MCMC so that it takes advantages of developments in high performance computing, and to improve MCMC performance through automated proposal construction. Following the algorithm, we began with a line-search step in which we iteratively swept through parameter space, one parameter at a time, calculating the posterior probability of each parameter set. Each line search swept through the parameter space many times, resulting in a "best" parameter set for each search. Line searches, however, generally get stuck at local maxima, or on ridges of the posterior probability surface. We therefore ran 2000 independent line-searches, with each started by randomizing the order of parameters in the search and the initial position in parameter space. Each search then produced a best parameter set, by maximizing the posterior probability.

Next, we used the top 5% of these 2000 parameter sets to construct automated proposal distributions for an MCMC routine. When MCMC is used to fit nonlinear models with more than a few parameters, correlations between parameters commonly disrupt the performance of the MCMC algorithm, by creating ridges on the surface of the posterior probability. To overcome this problem, we followed the line search-MCMC algorithm by using the top 5% of the 2000 parameter sets from the line searches, as ordered by decreasing posterior probability, in a PCA. We then used the PCA results to generate proposals for MCMC.

To explain how the PCA results were used, we note first that, in a standard Metropolis-Hastings MCMC algorithm, a random value of a single parameter is drawn from a proposal distribution, a new likelihood is calculated for the new parameter set, and then a decision is made to keep or reject the newly proposed parameter value. Following line search-MCMC, we instead drew a parameter in PCA space, and then we back-calculated to produce a new full set of model parameter values on the original, non-PCA-transformed scale. In each MCMC iteration, we therefore proposed a full, uncorrelated set of parameters, before calculating likelihoods. This vastly improved the mixing of our MCMC chains.

We ran 10 independent MCMC chains for each of our 1-$\bar{\nu}$ and 2-$\bar{\nu}$ models for at least 30,000 iterations, thinning by 100 iterations. This took approximately one week of computing or "wall-clock time" for each model. We tested for convergence via the Potential Scale Reduction Factor, $\hat{R}$, (Gelman et al., 2014), and we used visual diagnostics to assess overall mixing, and within- and among-chain auto-correlations and parameter correlations. Our results are based on strong convergence, except for models for which heterogeneity $C = 0$. As we describe in the main text, the latter models fit the data very poorly, so it may be that there is no Bayesian algorithm that will converge for those models.

#### 7.2 The Conceptual Basis of WAIC

The fundamental goal of model selection is to identify a model that will best predict future data (Konishi and Kitagawa, 2008). In a Bayesian context, the accuracy of the model predictions can be summarized by the posterior predictive density, the probability of future data given current data (Gelman et al., 2014);

$$p(\tilde{D}|D) \quad = \quad \int p(\tilde{D}|\theta)p(\theta|D)d\theta. \tag{A7}$$

Here $\tilde{D}$ is future data, $D$ is current data, and $\theta$ is again a vector of parameters. The predictive accuracy of the model $p(\tilde{D}|D)$ is thus calculated in terms of the probability of future data given the parameters $p(\tilde{D}|\theta)$, and in terms of the posterior distribution of the parameters $p(\theta|D)$. By definition we have not yet collected future data, so the ability of a model to predict future data must be approximated by the model's ability to predict existing data. The likelihood of current data, however, is an over-estimate of the likelihood of future data, because fitting a model to a current data set usually leads to the model being over-tuned to the current data (Konishi and Kitagawa, 2008). For this reason, raw likelihood scores are biased indicators of the best model.

Model selection criteria represent attempts to correct for this bias. In WAIC, the correction is calculated in terms of the variance of the posterior predictive density, which can be approximated by the variance of the likelihood, calculated using a sample from the posterior (Watanabe, 2009). The WAIC is then defined as (Gelman et al., 2014):

$$\text{WAIC} \quad = \quad \sum_{i=1}^{n} \log\left(\frac{1}{S}\sum_{s=1}^{S} p(D_i|\theta^s)\right) + \sum_{i=1}^{n} V_{s=1}^{S} \log p(D_i|\theta^s)$$

The first term, $\sum_{i=1}^{n} \log\left(\frac{1}{S}\sum_{s=1}^{S} p(D_i|\theta^s)\right)$, is the average likelihood across a large sample from the posterior. This average serves as an approximation to the expected log of the predictive density in equation (A7), a goodness of fit term. Here the likelihood is averaged over the $n$ data points (our 12 epizootic time series), over $S$ random parameter sets drawn from the posterior, and over the stochasticity in the model output.

The second term in equation (A8), $\sum_{i=1}^{n} V_{s=1}^{S} \log p(D_i|\theta^s)$, penalizes more complex models in terms of $V_{s=1}^{S}$, the variance of the log likelihood, again calculated across a large sample from the posterior. This term is an approximation to the variance of the log predictive density, here averaged over the $n$ data points and $S$ random draws, which inevitably increases with the number of parameters. The underlying idea is that more complex models will necessarily have better average log-likelihood scores $\sum_{i=1}^{n} \log\left(\frac{1}{S}\sum_{s=1}^{S} p(D_i|\theta^s)\right)$. Because more complex models are tuned to previous data, however, they often do a poor job of predicting future data. In such cases the more complex models will have higher average variance terms $\sum_{i=1}^{n} V_{s=1}^{S}$.

In order for the model with experiment-based priors to have a better WAIC score than the model with vague priors, the experimental priors must be located in areas of high likelihood scores within parameter space. In other words, our experiments have to have produced reasonably accurate estimates of the model parameters, which in turn requires that the processes in our experiments must roughly mimic processes in epizootics. In principle, there is no reason why this would not be the case. Nevertheless, because our transmission experiments were carried out at the scale of single branches, whereas the epizootic data were collected at the scale of entire forests, it seemed unlikely that the experiment-based priors would be centered in areas of high likelihoods.

Our goal in using WAIC was therefore to quantify the extent to which our experiment-based priors were located far enough from areas of high likelihood that one could reasonably argue that the model with vague priors provides a meaningfully better fit to the epizootic data, and thus that large-scale phenomena are helping to drive transmission. In making this determination, we follow the standard convention of requiring that models must differ in WAIC score by 3 or 4 in order for the better model to be considered as meaningfully better. Also, we included models that differed in which parameters had experiment-based versus vague priors. Because different parameters represent different processes, this allows us to test which processes were best captured by our experiments.

#### 7.3 Tests of WAIC

Exploratory simulations demonstrated that, when the number of draws of parameter sets from the posterior was more than about 500, further increases in the number of draws had only small effects on the WAIC score. In contrast, the number of realizations of the stochastic model had a strong effect on the WAIC score, apparently because the number of realizations has strong effects on the penalty term of the WAIC (fig. A6). Accordingly, for each model, we calculated WAIC scores based on 750 parameter sets drawn from the posteriors, and by estimating likelihoods ($\hat{\hat{L}}$) using 1000 realizations of each epizootic.

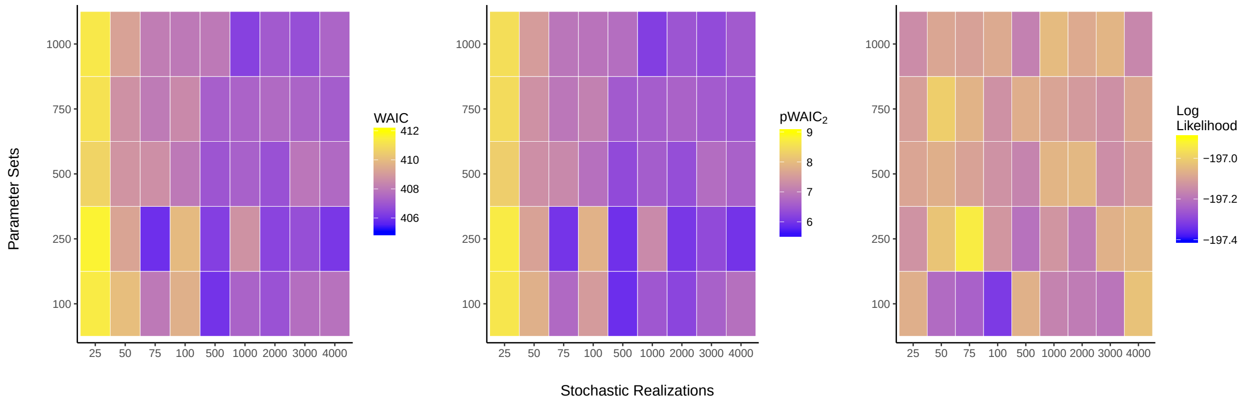

Figure A6: Effects of the number of parameter sets and the number of stochastic realizations on WAIC scores. Effects on the WAIC (left panel) are separated into effects on the penalty term (middle panel), and effects on the log likelihood (right panel). This simulation was carried out for the model that has an experiment-based prior on $\delta$, which has a WAIC score that is close to that of the best model.

#### 8 Visual fits to the observational data for models with intermediate numbers of experiment-based priors.

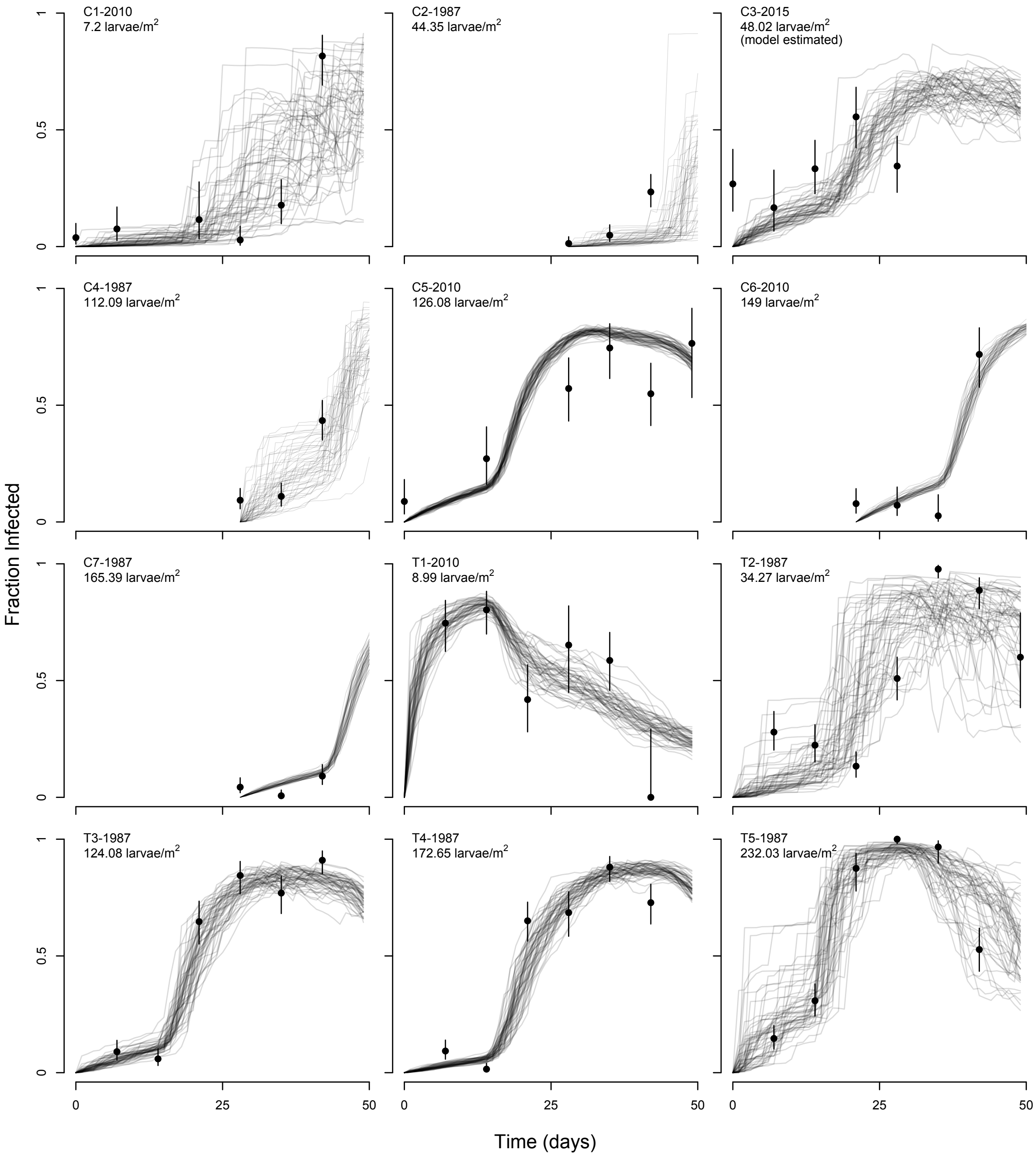

Figure A7: Stochastic realizations of the model that has experiment-based priors on the transmission heterogeneity parameter $C$, the ratio parameter $\rho$, and the death rate $\delta$.

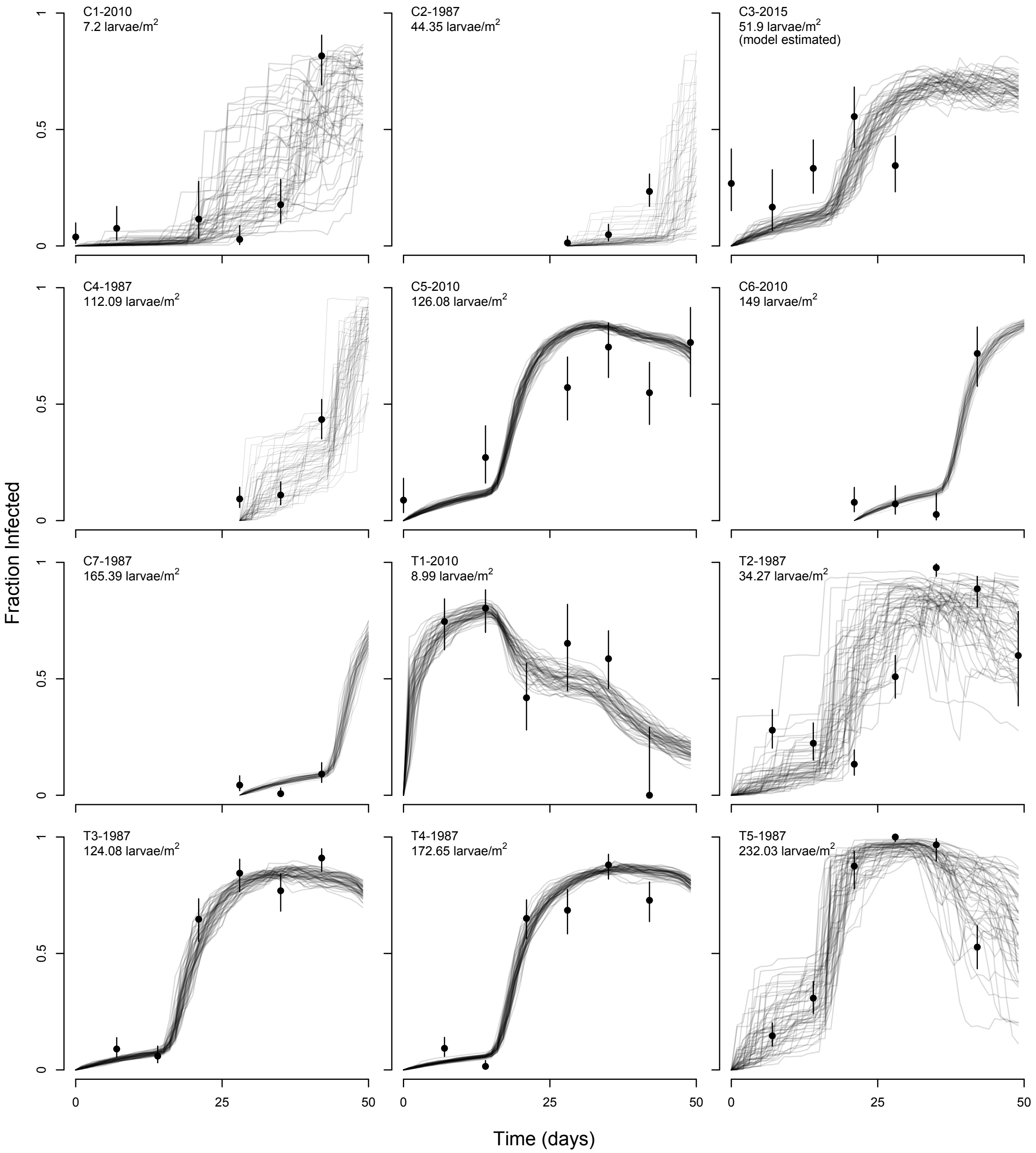

Figure A8: Stochastic realizations of the model that has an experiment-based prior on the transmission heterogeneity parameter $C$.

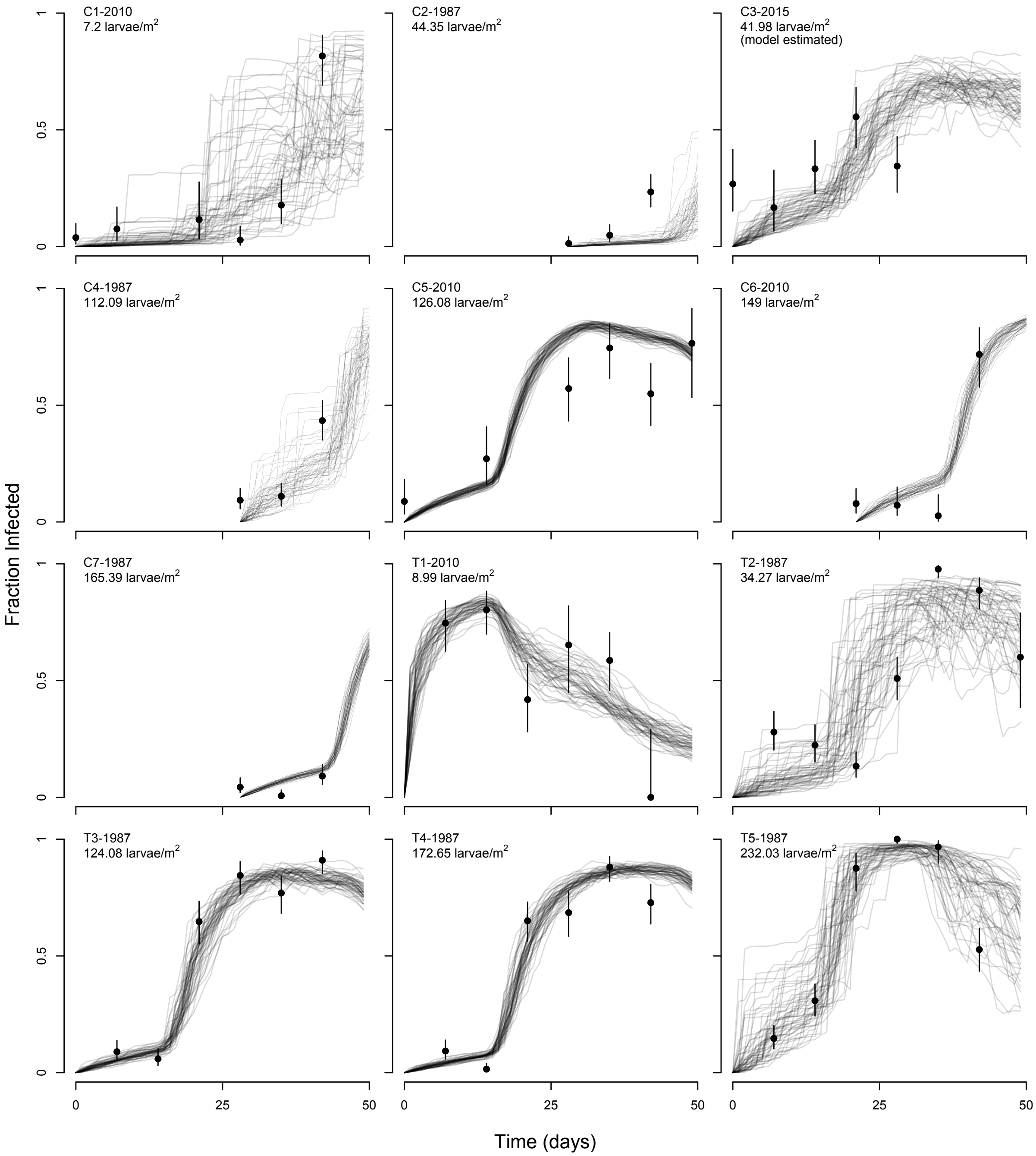

Figure A9: Stochastic realizations of the model that has an experiment-based prior on the ratio parameter $\rho$.

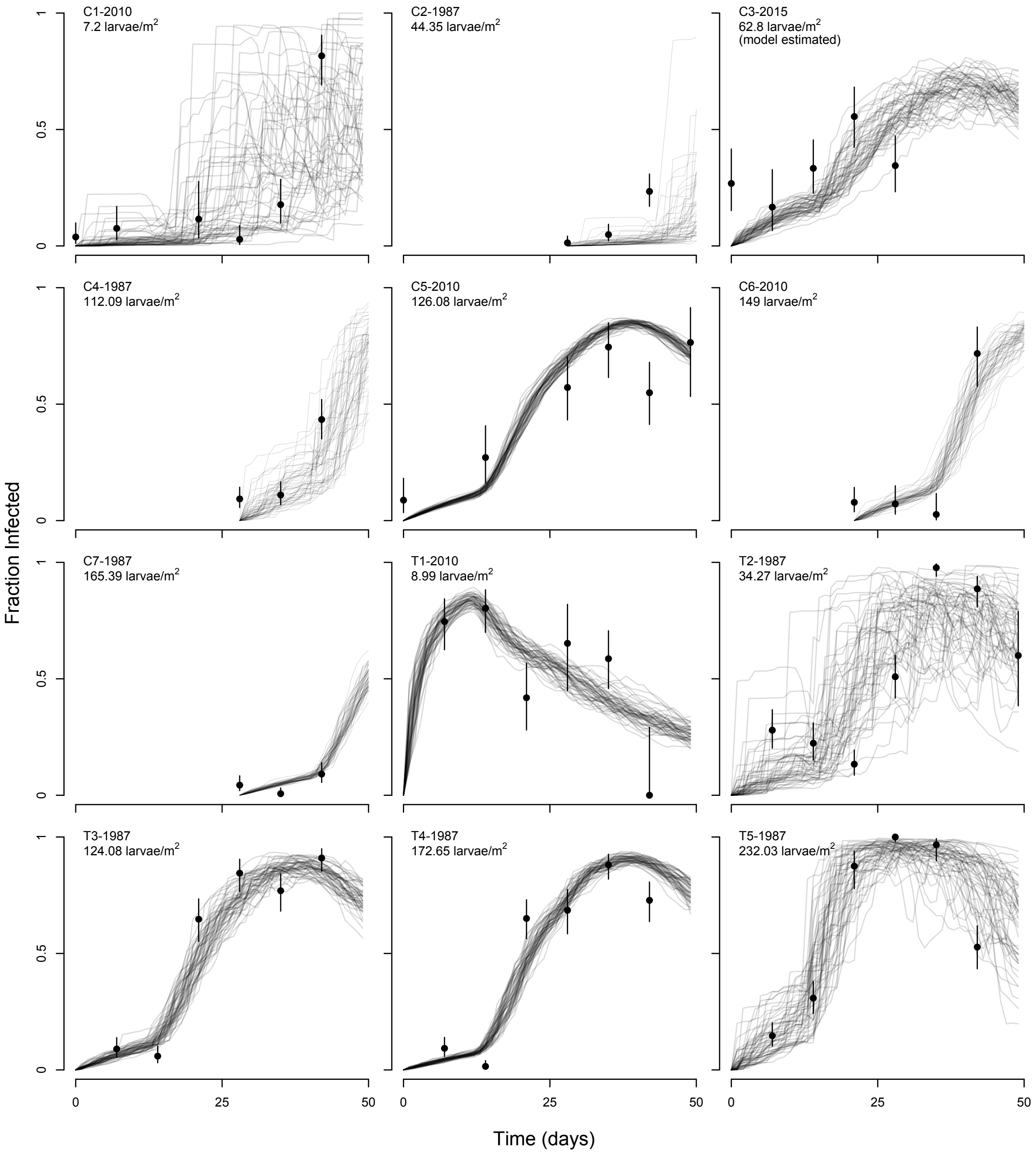

Figure A10: Stochastic realizations of the model that has an informative prior on the average transmission parameter $\bar{\nu}$.

### 9 Marginal posteriors all models

Here we present tables showing the marginal posterior parameter estimates for all of our models.

Table A2: Marginal posterior estimates of model parameters. The parameter $\gamma$ is an inverse measure of the extra-binomial variation in fraction infected, so that values of $\gamma < 1$ indicate strong over-dispersion. Given that in our case all values of $\gamma > 2$, overdispersion was moderate but not strong. Estimates are medians with 95% credible intervals. These are not on a log scale.

| Parameter | Model (Informative Priors) | Estimate |
|---|---|---|
| | $\delta$ | 0.061 (0.055, 0.067) |
| | None | 0.063 (0.056, 0.07) |
| | $C\ \rho\ \delta$ | 0.067 (0.059, 0.076) |
| $\delta$ | $C$ | 0.063 (0.058, 0.07) |
| | $C\ \rho\ \delta\ \bar{v}$ | 0.077 (0.066, 0.087) |
| | $\rho$ | 0.065 (0.057, 0.076) |
| | $\bar{v}$ | 0.083 (0.064, 0.103) |
| | $\delta$ | 1.360 (1.163, 1.588) |
| | None | 1.275 (1.108, 1.530) |
| | $C\ \rho\ \delta$ | 1.150 (0.968, 1.337) |
| $C$ | $C$ | 1.233 (1.024, 1.436) |
| | $C\ \rho\ \delta\ \bar{v}$ | 0.978 (0.813, 1.163) |
| | $\rho$ | 1.132 (0.963, 1.346) |
| | $\bar{v}$ | 0.836 (0.605, 1.086) |

| | | |
|---|---|---|
| | $\delta$ | 0.041 (0.001, 0.201) |
| | None | 0.042 (0.001, 0.236) |
| | $C\ \rho\ \delta$ | 0.016 (0, 0.117) |
| $\mu$ | $C$ | 0.053 (0.002, 0.221) |
| | $C\ \rho\ \delta\ \bar{v}$ | 0.007 (0, 0.037) |
| | $\rho$ | 0.031 (0.001, 0.255) |
| | $\bar{v}$ | 0.017 (0, 0.073) |
| | $\delta$ | 0.277 (0.082, 0.545) |
| | None | 0.166 (0.055, 0.377) |
| | $C\ \rho\ \delta$ | 0.059 (0.029, 0.101) |
| $\bar{v}$ | $C$ | 0.136 (0.053, 0.267) |
| | $C\ \rho\ \delta\ \bar{v}$ | 0.024 (0.014, 0.037) |
| | $\rho$ | 0.07 (0.025, 0.14) |
| | $\bar{v}$ | 0.017 (0.008, 0.027) |
| | $\delta$ | 0.004 (0.001, 0.012) |
| | None | 0.008 (0.002, 0.02) |
| | $C\ \rho\ \delta$ | 0.024 (0.01, 0.043) |
| $\rho$ | $C$ | 0.01 (0.002, 0.023) |
| | $C\ \rho\ \delta\ \bar{v}$ | 0.047 (0.02, 0.083) |
| | $\rho$ | 0.023 (0.008, 0.044) |
| | $\bar{v}$ | 0.062 (0.025, 0.123) |
| | $\delta$ | 2.117 (1.801, 2.426) |
| | None | 2.102 (1.831, 2.367) |
| | $C\ \rho\ \delta$ | 2.147 (1.864, 2.501) |
| $\gamma$ | $C$ | 2.156 (1.886, 2.456) |
| | $C\ \rho\ \delta\ \bar{v}$ | 2.248 (1.995, 2.531) |

| | |
|---|---|
| $\rho$ | 2.079 (1.704, 2.384) |
| $\bar{\nu}$ | 2.157 (1.899, 2.455) |

Table A3: Marginal posterior estimates of initial viral (i.e. infectious 1st instar cadaver) densities, and of initial host density for C3-2015. Estimates are medians with 95% credible intervals. These are not on a log scale.

| Parameter | Model (Informative Priors) | Estimate |
|---|---|---|
| | $\delta$ | 3.223 (1.314, 6.269) |
| | None | 4.259 (2.354, 8.446) |
| | $C\ \rho\ \delta$ | 8.272 (3.173, 17.101) |
| $P_0$, T1-2010 | $C$ | 4.099 (1.972, 7.311) |
| | $C\ \rho\ \delta\ \bar{\nu}$ | 13.271 (4.126, 30.747) |
| | $\rho$ | 7.38 (2.498, 13.666) |
| | $\bar{\nu}$ | 19.551 (6.467, 45.627) |
| | $\delta$ | 0.01 (0.001, 0.041) |
| | None | 0.02 (0, 0.105) |
| | $C\ \rho\ \delta$ | 0.052 (0.002, 0.216) |
| $P_0$, T2-1987 | $C$ | 0.029 (0, 0.159) |
| | $C\ \rho\ \delta\ \bar{\nu}$ | 0.152 (0.008, 0.698) |
| | $\rho$ | 0.065 (0.004, 0.477) |
| | $\bar{\nu}$ | 0.23 (0.005, 1.259) |
| | $\delta$ | 0.02 (0.004, 0.062) |
| | None | 0.045 (0.006, 0.173) |
| | $C\ \rho\ \delta$ | 0.127 (0.003, 0.437) |
| $P_0$, T3-1987 | $C$ | 0.054 (0.015, 0.121) |
| | $C\ \rho\ \delta\ \bar{\nu}$ | 0.333 (0.086, 0.793) |
| | $\rho$ | 0.115 (0.029, 0.286) |

| | | |
|---|---|---|
| | $\bar{v}$ | 0.538 (0.133, 1.227) |
| | $\delta$ | 0.018 (0.001, 0.057) |
| | None | 0.026 (0.001, 0.159) |
| | $C\ \rho\ \delta$ | 0.075 (0.019, 0.171) |
| $P_0$, T4-1987 | $C$ | 0.044 (0.012, 0.095) |
| | $C\ \rho\ \delta\ \bar{v}$ | 0.221 (0.053, 0.519) |
| | $\rho$ | 0.091 (0.018, 0.261) |
| | $\bar{v}$ | 0.374 (0.085, 0.889) |
| | $\delta$ | 0.062 (0, 0.315) |
| | None | 0.121 (0.027, 0.299) |
| | $C\ \rho\ \delta$ | 0.234 (0.014, 0.827) |
| $P_0$, T5-1987 | $C$ | 0.104 (0, 0.679) |
| | $C\ \rho\ \delta\ \bar{v}$ | 0.462 (0.072, 1.242) |
| | $\rho$ | 0.244 (0.031, 0.893) |
| | $\bar{v}$ | 0.724 (0.109, 2.1) |
| | $\delta$ | 47.855 (6.619, 148.114) |
| | None | 49.627 (7.389, 124.164) |
| | $C\ \rho\ \delta$ | 47.754 (5.464, 134.17) |
| $S_0$, C3-2015 | $C$ | 51.902 (2.506, 179.48) |
| | $C\ \rho\ \delta\ \bar{v}$ | 62.45 (4.44, 171.538) |
| | $\rho$ | 41.987 (3.418, 133.317) |
| | $\bar{v}$ | 62.803 (10.467, 146.299) |

Table A4: Marginal posterior estimates of transmission variability, $\sigma_{\epsilon_t}$, which is plot-specific. Estimates are medians with 95% credible intervals. These are not on a log scale.

| Plot | Model (Informative Priors) | Estimate |
|---|---|---|
| | $\delta$ | 2.742 (1.405, 5.048) |
| | None | 3.546 (1.246, 6.572) |
| | $C\ \rho\ \delta$ | 4.293 (2.698, 6.609) |
| C1-2010 | $C$ | 3.832 (1.893, 6.23) |
| | $C\ \rho\ \delta\ \bar{v}$ | 4.817 (3.393, 6.959) |
| | $\rho$ | 3.942 (2.033, 7.225) |
| | $\bar{v}$ | 5.641 (3.767, 7.924) |
| | $\delta$ | 3.537 (0.337, 12.439) |
| | None | 4.011 (2.165, 6.368) |
| | $C\ \rho\ \delta$ | 3.592 (0.758, 8.179) |
| C2-1987 | $C$ | 3.993 (0.74, 9.74) |
| | $C\ \rho\ \delta\ \bar{v}$ | 3.788 (0.646, 10.17) |
| | $\rho$ | 1.405 (0.132, 6.094) |
| | $\bar{v}$ | 2.705 (0.384, 7.888) |
| | $\delta$ | 0.769 (0.031, 3.865) |
| | None | 0.804 (0.006, 5.441) |
| | $C\ \rho\ \delta$ | 0.794 (0.02, 4.079) |
| C3-2015 | $C$ | 0.488 (0.001, 4.907) |
| | $C\ \rho\ \delta\ \bar{v}$ | 0.483 (0.001, 4.056) |
| | $\rho$ | 0.873 (0.02, 7.322) |
| | $\bar{v}$ | 0.5 (0.003, 3.693) |
| | $\delta$ | 3.636 (1.578, 6.299) |

C4-1987

| | | |
|---|---|---|
| | None | 2.597 (0.791, 6.158) |
| | $C\ \rho\ \delta$ | 1.959 (0.093, 9.11) |
| | $C$ | 2.858 (0.702, 6.928) |
| | $C\ \rho\ \delta\ \bar{v}$ | 1.86 (0.325, 5.442) |
| | $\rho$ | 1.535 (0.024, 6.543) |
| | $\bar{v}$ | 1.547 (0.048, 6.693) |
| | $\delta$ | 0.111 (0.002, 0.754) |
| | None | 0.117 (0.003, 0.775) |
| | $C\ \rho\ \delta$ | 0.099 (0, 0.866) |
| C5-2010 | $C$ | 0.084 (0, 1.019) |
| | $C\ \rho\ \delta\ \bar{v}$ | 0.097 (0.001, 0.974) |
| | $\rho$ | 0.087 (0.001, 0.962) |
| | $\bar{v}$ | 0.052 (0, 0.717) |
| | $\delta$ | 0.102 (0, 1.018) |
| | None | 0.154 (0.002, 1.216) |
| | $C\ \rho\ \delta$ | 0.125 (0.001, 1.437) |
| C6-2010 | $C$ | 0.098 (0.001, 0.914) |
| | $C\ \rho\ \delta\ \bar{v}$ | 0.128 (0, 1.453) |
| | $\rho$ | 0.202 (0.006, 1.162) |
| | $\bar{v}$ | 0.333 (0.012, 1.927) |
| | $\delta$ | 0.095 (0.001, 0.937) |
| | None | 0.222 (0.006, 1.261) |
| | $C\ \rho\ \delta$ | 0.098 (0, 1.443) |
| C7-1987 | $C$ | 0.152 (0.006, 0.92) |
| | $C\ \rho\ \delta\ \bar{v}$ | 0.12 (0.002, 1.119) |
| | $\rho$ | 0.106 (0.001, 0.724) |

| | | |
|---|---|---|
| | $\bar{v}$ | 0.136 (0.002, 0.796) |
| | $\delta$ | 0.368 (0.004, 2.743) |
| | None | 0.352 (0.012, 1.921) |
| | $C\ \rho\ \delta$ | 0.41 (0.012, 2.487) |
| T1-2010 | $C$ | 0.345 (0.002, 2.443) |
| | $C\ \rho\ \delta\ \bar{v}$ | 0.467 (0.003, 3.2) |
| | $\rho$ | 0.404 (0.003, 4.09) |
| | $\bar{v}$ | 0.21 (0, 2.63) |
| | $\delta$ | 2.976 (0.676, 7.66) |
| | None | 2.896 (0.997, 6.441) |
| | $C\ \rho\ \delta$ | 3.499 (1.576, 6.053) |
| T2-1987 | $C$ | 3.218 (0.166, 9.305) |
| | $C\ \rho\ \delta\ \bar{v}$ | 3.093 (1.504, 5.387) |
| | $\rho$ | 2.534 (0.829, 6.221) |
| | $\bar{v}$ | 2.92 (1.58, 4.817) |
| | $\delta$ | 0.294 (0.003, 2.722) |
| | None | 0.237 (0.003, 3.072) |
| | $C\ \rho\ \delta$ | 0.571 (0.003, 4.353) |
| T3-1987 | $C$ | 0.223 (0.001, 2.573) |
| | $C\ \rho\ \delta\ \bar{v}$ | 0.432 (0.002, 2.975) |
| | $\rho$ | 0.35 (0, 2.723) |
| | $\bar{v}$ | 0.359 (0.002, 2.113) |
| | $\delta$ | 0.105 (0, 1.598) |
| | None | 0.105 (0.001, 1.194) |
| | $C\ \rho\ \delta$ | 0.363 (0.007, 2.03) |
| T4-1987 | $C$ | 0.118 (0.001, 1.002) |

| | | |
|---|---|---|
| | $C\ \rho\ \delta\ \bar{v}$ | 0.265 (0.001, 2.258) |
| | $\rho$ | 0.21 (0.004, 1.618) |
| | $\bar{v}$ | 0.16 (0.001, 1.936) |
| | $\delta$ | 2.411 (0.008, 14.717) |
| | None | 2.397 (0.066, 8.573) |
| | $C\ \rho\ \delta$ | 2.752 (0.15, 10.398) |
| T5-1987 | $C$ | 2.966 (0.557, 7.488) |
| | $C\ \rho\ \delta\ \bar{v}$ | 2.56 (0.579, 6.598) |
| | $\rho$ | 2.325 (0.24, 6.59) |
| | $\bar{v}$ | 1.818 (0.071, 8.203) |

### 10 JAGS model statement for fitting the transmission model to experimental data

```
model{

##########
# Priors #
##########

  # Virus-level transmission parameters.
  # Three viruses: TMB-1, WA, NM

  for(v in 1:3){
    # C and nu.bar (log scale):
    nC[v] ~ dnorm(0, 1 / (1 ^ 2))
    nnu.bar[v] ~ dnorm(-8, 1 / (2 ^ 2))

    C[v] <- exp(nC[v])
    nu.bar[v] <- exp(nnu.bar[v])
  }

  # Error in Initial Cadaver Density:
  # Error in cadaver density varies by experimental treatment.
  # (i.e. 1, 2, 3 represent 0, 10, or 40 initial cadavers)

  for(d in 1:3){
    p.mean[d] ~ dnorm(0, .001)
```

```
    p.sd[d] ~ dgamma(0.001,0.001)

    p.tau[d] <- 1 / (p.sd[d])^2

  }

  # Ratio parameter:

  # Informative prior generated by counting virus in

  # first and fourth instar larvae

  nratio ~ dnorm(-3.387, 1 / 0.25^2)

  ratio <- exp(nratio)

##############

# LIKELIHOOD #

##############

  for(i in 1:N.obs){

    # Binomial likelihood:

    # n.inf = number infected from each bag

    # n.recov = total number recovered from each bag at end of experiment

    n.inf[i] ~ dbin(prob[i], n.recov[i])

    # Heterogeneity model:

    prob[i] <- 1 - (1 + (C[Virus[i]])^2 * nu.bar[Virus[i]] *

    (P[i]+0.00001) * ratio * 7)^(-1/(C[Virus[i]]^2))

  }
```

```
########################
# ERROR IN CAD. DENSITY #
########################
  # P = estimated cadaver density,
  # based on estimated area of foliage in each bag
  # Cad.Den = vector of treatment identifiers
  # (i.e. 1, 2, 3 represent 0, 10, or 40 initial cadavers)

  for(i in 1:N.obs){
    P[i] ~ dnorm(p.mean[Cad.Den[i]], p.tau[Cad.Den[i]])
  }

}
```

### 11 JAGS model statement to fit a Gamma distribution to the time-to-death data

```
model{

##########
# Priors #
##########

  g_alpha ~ dunif(0, 100)
  g_beta ~ dunif(0, 100)

##############
# LIKELIHOOD #
##############
  # y = vector with time-to-death for each larva

  for(i in 1:n_samp){
    y[i] ~ dgamma(g_alpha, g_beta)
  }

}
```

### Literature Cited

Bolker, B. M. 2008. Ecological models and data in R. Princeton University Press.

D'Amico, V., J. S. Elkinton, J. D. Podgwaite, J. Buonaccorsi, and G. Dwyer. 2005. Pathogen clumping: an explanation for non-linear transmission of an insect virus. Ecological Entomology 30:383–390.

Dwyer, G. 1991. The effects of density, stage and spatial heterogeneity on the transmission of an insect virus. Ecology 72:559–574.

Dwyer, G., J. Firestone, and T. E. Stevens. 2005. Should models of disease dynamics in herbivorous insects include the effects of variability in host-plant foliage quality? American Naturalist 165:16–31.

Eakin, L., M. Wang, and G. Dwyer. 2015. The effects of the avoidance of infectious hosts on infection risk in an insect-pathogen interaction. Am. Nat. 185:pp. 100–112.

Gelman, A., J. B. Carlin, H. S. Stern, D. B. Dunson, A. Vehtari, and D. B. Rubin. 2014. Bayesian Data Analysis, Third Edition. Chapman & Hall/CRC Press. New York, NY.

Han, X., and P. E. Kloeden. 2017. Random Ordinary Differential Equations and Their Numerical Solution. Springer.

Kennedy, D. A., V. Dukic, and G. Dwyer. 2014. Pathogen Growth in Insect Hosts: Inferring the Importance of Different Mechanisms Using Stochastic Models and Response-Time Data. American Naturalist 184:407–423.

Kernighan, B. W., and D. M. Ritchie. 2006. The C programming language.

Konishi, S., and G. Kitagawa. 2008. Information criteria and statistical modeling. Springer Science & Business Media.

Mason, R., D. Scott, and H. Paul. 1993. Forecasting Outbreaks of the Douglas-Fir Tussock Moth From Lower Crown Cocoon Samples. USDA Forest Service Research Paper PNW-RP-460.

Mason, R., and T. Torgersen. 1983. Mortality of larvae in stocked cohorts of the Douglas-fir tussock moth, *Orgyia pseudotsugata* (Lepidoptera: Lymantriidae). Canadian Entomologist 115:1119–1127.

Mason, R. R., R. Beckwith, and H. G. Paul. 1977. Fecundity Reduction During Collapse of a Douglas-fir Tussock Moth Outbreak in Northeast Oregon. Environmental Entomology 6:623–626.

Otvos, I. S., J. C. Cunningham, and L. M. Friskie. 1987. Aerial application of nuclear polyhedrosis virus against douglas-fir tussock moth, *Orgyia pseudostugata* (Mcdunnough) (Lepidoptera: Lymantriidae). 1. impact in the year of application. Canadian Entomologist 119:697–706.

Parker, B. J., B. D. Elderd, and G. Dwyer. 2010. Host behaviour and exposure risk in an insect-pathogen interaction. Journal of Animal Ecology 79:863–870.

Polivka, K., G. Dwyer, K. C. Skalisky, C. J. Mehmel, and J. L. Novak. 2012. Analysis of NPV epizootics during a Douglas-fir tussock moth management project in the Methow Ranger District (Okanogan-Wenatchee National Forest), 2010. Report to Okanogan-Wenatchee National Forest, Forest Health Protection .

Scott, D., and L. Spiegel. 2002. One and two year follow-up evaluation of TM Biocontrol-1 treatments to suppress Douglas-fir tussock moth in the Blue Mountains of northeastern Oregon and southeastern Washington. Technical Report BMPMSC-02-02, USDA Forest Service, Pacific Northwest Region.

Watanabe, S. 2009. Algebraic geometry and statistical learning theory, vol. 25. Cambridge University Press.
